# Operator model for evolutionary dynamics

**DOI:** 10.1101/2023.11.12.566730

**Authors:** Kangbien Park, Yonghee Bae

## Abstract

Drift, selection, and mutation are integral evolutionary factors. In this article, *operator model* is newly suggested to intuitively represent those evolutionary factors into mathematical operators, and to ultimately offer unconventional methodology for understanding evolutionary dynamics. To be specific, each of the drift, selection, and mutation was respectively interpreted as operator which in essence is a random matrix that acts upon the vector which contains population distribution information. The simulation results from the operator model coincided with the previous theoretical results for beneficial mutation accumulation rate in concurrent and successional regimes for asexually reproducing case. Furthermore, beneficial mutation accumulation in strong drift regime for asexually reproducing case was observed from the simulation while allowing the interactions of mutations with diverse selection coefficients. Lastly, methods to justify, reinforce, apply, and expand the operator model were discussed to scrutinize the implications of the model. With its unique characteristics, the operator model is expected to broaden perspective and to offer effective methodology for understanding the evolutionary process.

## 1. Introduction

Biological system faces evolution mainly through drift, selection, and mutation (Baake and Gabriel, 2000; Garcia-Dorado et al, 2007; Walsh and Lynch, 2018). When new mutations are provided, those mutations face selective pressure. Most of the case, the provided mutations are deleterious or neutral and do not offer advantages during selection. However, some of the mutations are beneficial for survival and when they fix, these mutations produce their own lineages (Levy et al, 2015; Nguyen Ba et al, 2019). If these lineages establish in the population, the lineages often interact with each other, which is found to be an important contributor for evolutionary process (Desai et al, 2007). Moreover, the random drift comes into play during the process of evolution and plays significant role in shaping the evolutionary facets (Ewens, 2004).

For describing the evolutionary process in quantitative terms, methods based on solving the master equation for evolution have steadily been used (Aalto, 1989; Ewens, 2004; De Oliveira, 2014). Among those methods, Wright-Fisher (Wright, 1949) model utilizes the Brownian diffusion model and solves the stochastic second order PDE which is given from the approximation of the probability density function (Ewens, 2004; Wakeley, 2005; Tran et al, 2013; Bräutigam and Smerlak, 2022). Moreover, Moran process (Moran, 1958; Ewens, 2004; Murihead, 2009) model assumes that birth rate is proportional to relative fitness, and if an offspring is born, the pre-existing individual should demise to maintain the total population number, which would eventually depict the evolutionary situation.

Whereas those models mainly focus on solving the master equation for random variables, the newly proposed *operator model* in this article focuses on the mathematical operator which is essentially a random matrix (i.e., matrix with random variables for its entries) that contain the information for evolutionary process. With such attempt, the objective of operator modelling is to suggest a new methodology for tracking evolutionary dynamics where each of the evolutionary factor (i.e., drift, selection, mutation) could be intuitively represented in the form of operator, and to ultimately offer broader perspective for understanding the evolution process. As discrete-time Markov process, the analysis is proceeded with population vector |β (t) ⟩ which contains the population distribution information for each discrete generation t. To be specific, by acting logos operator 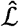 which is a random matrix that is constructed from the information of |β (t) ⟩ and newly defined tensors S and E, the information for next population could be explicitly found in the relation of 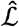 · |β (t) ⟩ = |β (t + 1) ⟩. Such operator representation is intuitive as it depicts a picture where evolutionary force 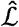 directly acts upon the population |β (t) ⟩ and affects its dynamical state. With this approach, analysis and simulation for general N (total population number), s (fitness value), and m (mutation rate) could efficiently be made where N may vary throughout the generation, and different s and m values may coexist in one generation. Moreover, the operator modelling could simply add a well-defined operator (e.g., sexual reproduction operator) to deal with any new evolutionary factors. Lastly, observation for any specific trait of interest throughout the generation could be effectively made.

## 2. Theory

Prior to any detailed analysis, basic conventions, definitions, and assumptions would be introduced. First, the vectors, matrices, and tensors for the analysis in this article generally have infinite number of elements to reflect the infinite possibility of evolution. However, since the population for generation t maintain finite number of existing mutations due to clonal interference when there are lot of mutations, only finite number of elements are of interest for vectors, matrices, and tensors. To reflect such characteristics and to minimize the total amount of consideration, the omission rule was defined. By this rule, 0 elements for vectors, identity parts for matrix, and 0 indices for tensors are *not represented*, although they exist.

Moreover, the basic assumption underlying in this theory is that the population faces mutation, selection, and drift in sequence. To represent such premise, generation t is divided into five parts of t_i_, t_m_, t_v_, t_d_, t_f_. Here, t_m_, t_v_, t_d_ each signifies the period when the population suffers mutation, selection, and drift in generation t. Moreover, t_i_ signifies the initial stage (before mutation) and t_f_ signifies the final stage (after drift). Finally, when the generation is expressed as t (without lower index), it conveys the general meaning of generation t and could correspond to all of the t_i_, t_m_, t_v_, t_d_, t_f_, depending on the context.

Now, the population set with corresponding population vector, and spanning and environment tensors should be defined. The population set *β*(t) is a set of population states {β ^1^(t), .., β^k^(t), .., β^n^(t)} where each β^1^(t), .., β^k^(t), .., β^n^(t) is referred to as the k^th^ “state” (k ∈ [1,*n* (*t*)]) (n(t) is the total number of trait for generation t) of generation t, and each of the state is a distinctive set of trait element. For detail, assume two trait sets f (color), g (shape) are the subject for evolutionary dynamics analysis. Each of the trait set f, g is consisted of the trait elements. Specifically, let there be blue (f_1_), red (f_2_) traits for color trait f (i.e., f = {f_1_, f_2_}), and sphere (g_1_), square (g_2_) traits for shape trait g (i.e., g = {g_1_, g_2_}) for generation T. Then, each state β^k^(T) (k ∈ [1,*n* (T)]) in set β(T) is composed with one of the trait elements in sets f, g thus yielding n(T) ≤ 2 × 2. For instance, let some state element β ^k^(T) in β(T) be {f_1_, g_2_}, thus having blue and square trait. For the rest of the analysis, β^k^(T) is defined to be {f_1_, g_2_}.

The rate of any specific mutation depends on the particular state β^k^(T) which it occurs from, as well as the environment that β^k^(T) is in (Ralston, 2008; Tan et al, 2009; Krašovec et al, 2014, Zhang, 2023). To represent such information, the newly defined spanning tensor S would be introduced. The spanning tensor 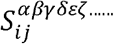 is essentially the library of the mutation rates and has infinite number of upper indices. The upper indices provide information regarding from which of the state the mutation is occurring, and lower indices represent which of the new mutation is of interest. Construction of spanning tensor, particularly the number of lower indices of the tensor could vary depending on the objective of analysis. For β^k^(T)= {f_1_, g_2_} analysis, 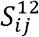 would allow the value of any particular mutation rates. For the rest of the analysis, let the mutation for k state in T generation occur for only f and g set with f_3_, f_4_, g_3_, g_4_ traits possibly added. Specifics about spanning tensor is introduced in the supplementary.

After the mutation step, the population faces a selective pressure, which is expressed with the newly defined environment tensor *E*^*αβγδεζ* … …^. The environment tensor is expected to be generated from theoretical fitness landscape (Gavrilets, 2010; de Visser and Krug, 2014), or through statistical analysis of experimental data. With the above example of β(T), the state β^k^(T) has its own designated fitness. That is, the trait f_1_, g_2_ in total designates the fitness for β^k^(T_v_), and the fitness is given as *r*^*k*^ = *E*^12^ for {f_1_, g_2_}. The *r*^*k*^ signifies relative fitness of kth state compared to the initial state of *r*^1^ = 1.

With these defined vectors and tensors, two equivalent representations in different forms, the “set representation” and “whole representation” could be constructed. Set representation, which offers concise expression for the computation process, is explained rigorously in the supplementary material. Here, the focus is on whole representation, which would give intuitions on how the operator modelling works.

The whole representation is based on the omission rule convention. For the β^k^(T) states in β(T) set, define a map *ψ*such that *ψ* (β^k^(T)) = x^k^ where x^k^ is the frequency of the state k. Also, define |β (t) ⟩ ≡ (*ψ* (β^1^(T)), …., *ψ* (β^k^(T)), .., *ψ* (β^n^(T))^T^ where 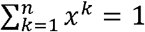. Here, note that |β (T) ⟩ ≡ (*ψ* (β^1^(T)), …., *ψ* (β^k^(T)), .., *ψ* (β^n^(T),0,0, ..)^T^ is a vector with infinite elements, but by omission rule, only the integral parts for the analysis is represented.

For the mutation, β^i, prior^(T_m_) should be considered for every i∈[1, n(T)]. Let the kth state β^k^(T_m_)= {f_1_, g_2_} be the subject to be considered. Then, β (T_m_) ⟩ = (ψ (β^1^(T_m_)), …., ψ (β^n^(T_m_)), .., *ψ* (β^n^(T)), *ψ* ({f_3_, g_2_}^k, prior^), *ψ* ({f_4_, g_2_}^k, prior^), *ψ* ({f_1_, g_3_}^k, prior^), *ψ* ({f_1_, g_4_}^k, prior^))^T^. The total number of represented prior sets in this example would be 2 × 2 = 4.

Using this system, the mutation operator 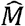 is essentially a random matrix containing the spanning tensor information represented as 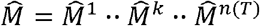 and acts on the vector 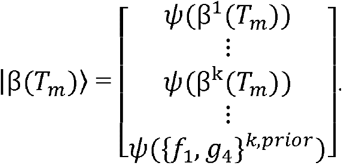. Each 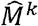 has the form of (1) or identity matrix.

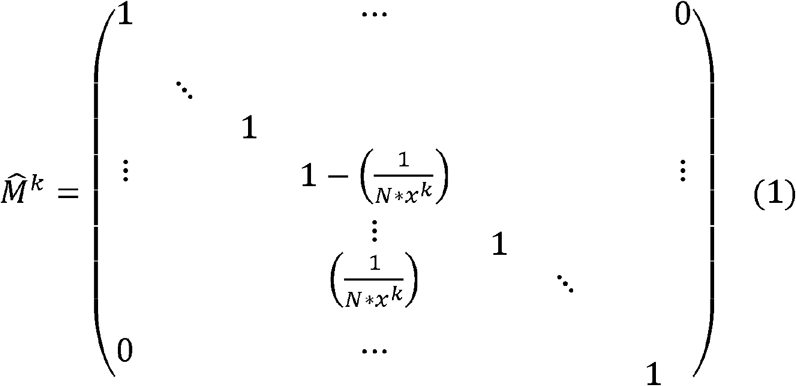

The probability for (1) form is given as 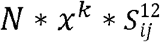, which is deduced from the expectation value of Binomial distribution with *N* * *x*^*k*^ for trial number and 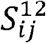 for probability of occurrence, while the probability for identity matrix is given as 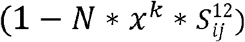. When 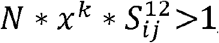, it loses the meaning of probability, and could be interpreted as 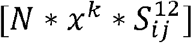 mutations certainly happening, and 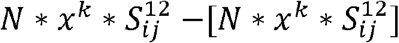 is interpreted as probability for extra mutation. In such case, (1) would have 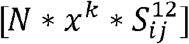 off-diagonal terms for certain, and may have 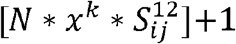 off-diagonal terms with probability 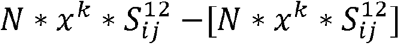. Further justification of such interpretation is dealt in discussion. For now, the following explanation assumes a situation where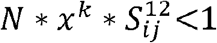 and 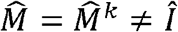 (i.e., single mutation only occurred for kth element). Table 1 shows the shape of mutation operator by allocating the indices for the 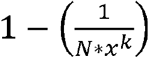 element and 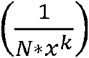 element.

**Table 1.** Random matrix and probability operating on β^k, prior^(T). For the application, multiple-fold mutations from one state are often unlikely to happen and such probabilities may be ignored.

| Probability | $\hat{M}_{whole}$ |
| --- | --- |
| $N \times x^k \times S_{32}^{12}$ | $\hat{M}_{k,k} = 1 - \left(\frac{1}{N * x^k}\right), \hat{M}_{k,n+1} = \left(\frac{1}{N * x^k}\right)$ |
| $N \times x^k \times S_{42}^{12}$ | $\hat{M}_{k,k} = 1 - \left(\frac{1}{N * x^k}\right), \hat{M}_{k,n+2} = \left(\frac{1}{N * x^k}\right)$ |
| $N \times x^k \times S_{13}^{12}$ | $\hat{M}_{k,k} = 1 - \left(\frac{1}{N * x^k}\right), \hat{M}_{k,n+3} = \left(\frac{1}{N * x^k}\right)$ |
| ... | ... |
| $N^2 \times (x^k)^2 \times S_{42}^{12} \times S_{14}^{12}$ | $\hat{M}_{k,k} = 1 - 2 * \left(\frac{1}{N * x^k}\right), \hat{M}_{k,n+2} = \left(\frac{1}{N * x^k}\right), \hat{M}_{k,n+4} = \left(\frac{1}{N * x^k}\right)$ |
| ... | ... |
| $1 - \text{sum of all the previous probability values}$ | $\mathbf{1}_{(n+4) \times (n+4)}$ |

As this example for the whole representation illustrates, consideration of possible mutations from one β^k^(T_m_) state yielded four additional prior terms. Therefore, to consider for the general cases where mutation happens for various states including the kth state, totally 4×n additional prior terms should be dealt with in this example since all the states of k∈[1, n(T)] respectively has four prior terms. If the number of state increases as generation passes, the mutation operator in the whole representation analysis may become too big and unwieldly to utilize for the analysis. In this case, the priorly introduced set representations would be handy to use with.

Now, the potential operator, where all the r values are defined from E tensor is expressed as (2)

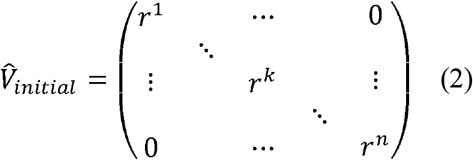

that acts on 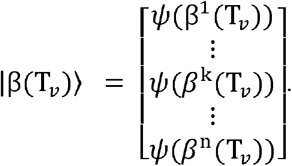.

However, the resulting vector 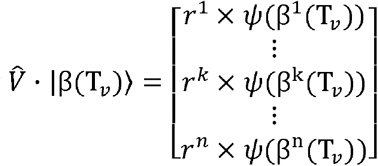 may not add up to 1.

Therefore, the additional step of dividing 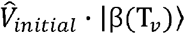 with 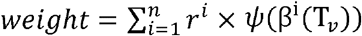 should be conducted to preserve the 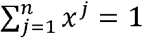 condition of |β (T) ⟩ and such weight information should be included in the final form of 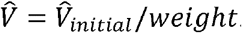. Hence, after the selection step, the whole representation may yield 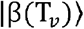.

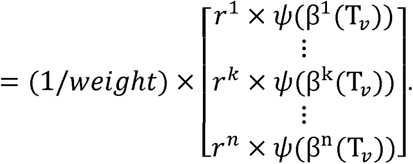

After the selection stage, the drift phenomenon is set to play important role for determining the evolutionary facets of population. The drift operator may be constructed based on various distribution, but in this model, the hypergeometric function was used to explain drift (Chesson, 1976; Fog, 2005; Fog 2008). Specifically, let |N ⟩_j_ = [*N* × |β (T_d_) ⟩_*j*_] where [] is a gauss function for ∀*j* ∈ [1, *n*(*T*)]. Then, |N_*initial*_⟩ = (*i*^1^, *i*^2^,.., *i*^*k*^,.., *i*^*n*^) where *i*^*k*^ = [N × ψ(β^k^(T))]. Now, assume that the drift is modeled by the situation where every state of β(T) duplicates its numbers to 2 × |N_*initial*_⟩ and only part of them survive after the drift with the numbers |N_*ultimate*_⟩ = (*u*^1^, *u*^2^,.., *u*^*k*^,.., *u*^*n*^,) which suffice the relation that 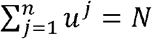 which is the environmental capacity value. This is where multivariate hypergeometric distribution comes in. The probability for |N_*ultimate*_⟩ from 2 × |N_*initial*_⟩ could be found with the multivariate hypergeometric distribution as

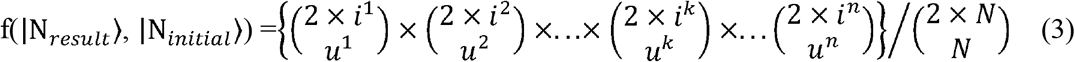

where f is the multivariate hypergeometric distribution. Here, 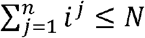, but since N is usually very large, assume that 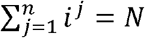 to set 2 × *N* value for 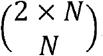.

Setting (*u*^*k*^/*N*) ≡*d*^*k*^,

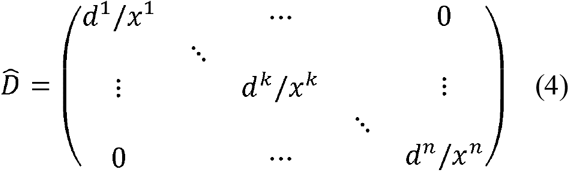

is the drift operator which acts on the total population vector 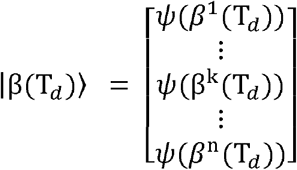. Here, 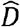 is essentially a random matrix where

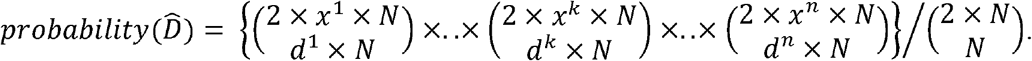

Finally, after the drift step, the big representation may yield 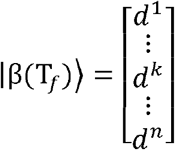 as a final form of generation T.

For the drift operator, strength of the drift could be determined by the coefficient that is multiplied in front of *i* in (3). The assumption was that the organism doubles for one generation and thus coefficient multiplied to *i* was 2, but such coefficient could vary depending on the situation. To express stronger drift, the coefficient multiplied to *i* could set to be larger.

Summing up all the operators intorduced, the logos operator 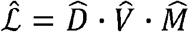 would have the form of ∞ × ∞ square matrix. Nevertheless, by the omission rule, the 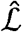 is represented with operators that satisfy 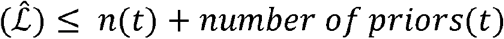 that act on β(t).

Moreover, from 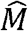, the 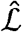 may have off-diagonal element by some probability given from the spanning tensor. From the logos operator of whole representation,

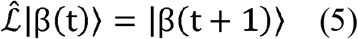

and information of β(t+1) could thus be derived from β(t).

## 3. Simulation

To decrease the complexity of computation, set representation was used instead of whole representation for the simulation. The aim of the simulation was to validate that operator model could reproduce and assess the pre-existing research results, and moreover understand the evolutionary dynamics in the high drift situation. To achieve this, rate of beneficial mutation accumulation in concurrent, successional regime was computed on the assumption that 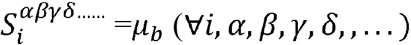, and *E*^*αβγδ……*^ = s × (number of nonzero indices *α, β, γ, δ, …*.) where *μ*_*b*_ and s are fixed parameters. For concurrent regime where clonal interference takes place (Gerrish and Lenski, 1998; Miralles et al, 1999; Park and Krugh, 2007), beneficial mutation accumulation rate was set to be v ≈ *s*^2^ [(2*ln* (*Ns*) − *ln* (*s*/ *μ*_*b*_))/ *ln*^2^ (*s*/ *μ*_*b*_)] (Desai et al, 2007), and fixation probability was set to be 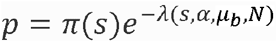 where *π* (*s*) ≈ 4*s* and 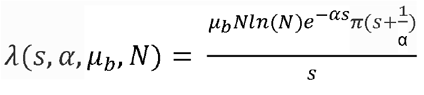 (Gerrish and Lenski, 1998). Here, α is a parameter that is determined from experiment or theory. Through simulation, fixation probability was measured for concurrent regime and α was predicted. For the successional regime, beneficial mutation accumulation rate was set to be v = N * *μ*_*b*_ * *s*^2^ (Desai et al, 2007). The specific parameters for concurrent regime were N between 10^5^ to 10^8^, m between 2 × 10^−8^ to 2 × 10^−5^, s between 0.005 to 0.003 and for successional regime were N between 10^3^ to 10^4^, m between 2 × 10^−7^ to 8 × 10^−5^, s between 0.001 to 0.003.

For the regime where drift is strong, beneficial mutation accumulation was observed for mutations with diverse s values drawn from Cauchy distribution 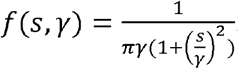. The shape of the distribution for s as Cauchy distribution was constructed from a crude approximation for previous research result (Böndel et al, 2022). Here, 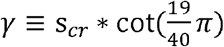 to let new mutation have s value between [-*s*_*cr*_, *s*_*cr*_] with the probability of 95% from the characteristics of Cauchy distribution. From such setting, if any new mutation was provided, the mutation rate was given from 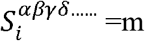 · probability value from Cauchy distribution for new trait, and to consider the accumulation of mutation, *E*^*αβγδ……*^ = selection coefficient for new trait+ parental *E*^*αβγδ……*^.

For each case of (N, m, s) parameters, with the fixed s values, 300 iterations were made to measure the proportion of simulated value/theoretical value for beneficial mutation accumulation rate in concurrent and successional regimes. Moreover, for each case of (N, m, s) parameters, with s values sampled from Cauchy distribution, 100 (generally done) or 300 (for more precise measurement) iterations were made for the strong drift regime to measure the average rate of beneficial mutation accumulation.

## 4. Results & Discussion

The 300 proportion values of (operator modelling simulation result/previous theoretical result) for beneficial mutation accumulation rate yielded mean values mostly in the range of [0.8, 1.2] for diverse parameters in concurrent and successional regimes which highly suggest that pre-exiting theories and operator modelling renders coinciding results. Moreover, the alpha value for fixation probability from Gerrish and Lenski’s formula (1998) was estimated as in the table 2 (more in supplementary). The muller plots in Fig. 1. from the operator modelling shows that as s value increases, the generation width of any fixed state increases, while as m value increases, the generation width of any fixed state decreases and variety for states increases. This is easily understood since new mutation has higher probability of fixation for larger s, and new mutations, which are the candidates for fixation, are more provided for larger m. Furthermore, the observed pattern of decreasing mean value for increasing s value could be observed in the data. Such phenomenon is predicted to happen not because of any implications within the operator modelling, but due to the approximation which previous theories were built upon. Such approximation was setting r^0^ =1, r^n^=(1+n × s), r^n^=(1+(n+1) × s), to imply that 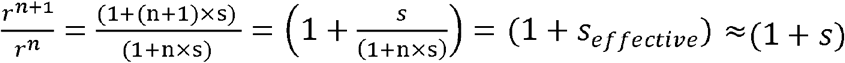, (here n means n-tuple mutation) which basically assumes that n × s is comparably smaller than 1. However, as s becomes larger such approximation loses accuracy and *s*_*effective*_ would decrease as (1+n × s) > 1, which then explains the decreasing behavior of the mean value. Hence, the simulated result from operator model well coincides with the pre-existing theories of evolutionary dynamics.

**Table 2.** Parameters and description.

| Parameters | Description |
| --- | --- |
| N | Total population number |
| $\mu_b$ | Rate for Beneficial mutation |
| m | General mutation rate including beneficial, neutral, and deleterious mutations |
| r | Relative fitness from initial trait |
| s | Selection coefficient, $s = r - 1$ |
| $s_{cr}$ | Limiting factor for s sampling from Cauchy distribution |
| v | Beneficial mutation accumulation rate |

**Table 3.** Part of the data for concurrent regime. The parameters are representing N, m, s values in order. The data measured mean and standard deviation of 300 proportion values of (operator modelling simulation result/previous theoretical result) for beneficial mutation accumulation rate. Moreover, alpha value for the fixation probability was estimated with the simulation.

| parameters | mean | stdev | estimated alpha |
| --- | --- | --- | --- |
| 1E+07, 2E-07, 0.005 | 0.9841 | 0.1115 | 729 |
| 1E+07, 2E-07, 0.02 | 0.9765 | 0.1332 | 181 |
| 1E+07, 2E-08, 0.005 | 1.0637 | 0.2078 | 412 |
| 1E+07, 2E-08, 0.02 | 0.9775 | 0.1197 | 100 |
| 1E+08, 2E-07, 0.005 | 0.9386 | 0.1110 | 1121 |
| 1E+08, 2E-07, 0.02 | 0.8428 | 0.0508 | 278 |
| 1E+08, 2E-08, 0.005 | 0.9887 | 0.1724 | 747 |
| 1E+08, 2E-08, 0.02 | 0.9097 | 0.0538 | 185 |

**Fig. 1.**
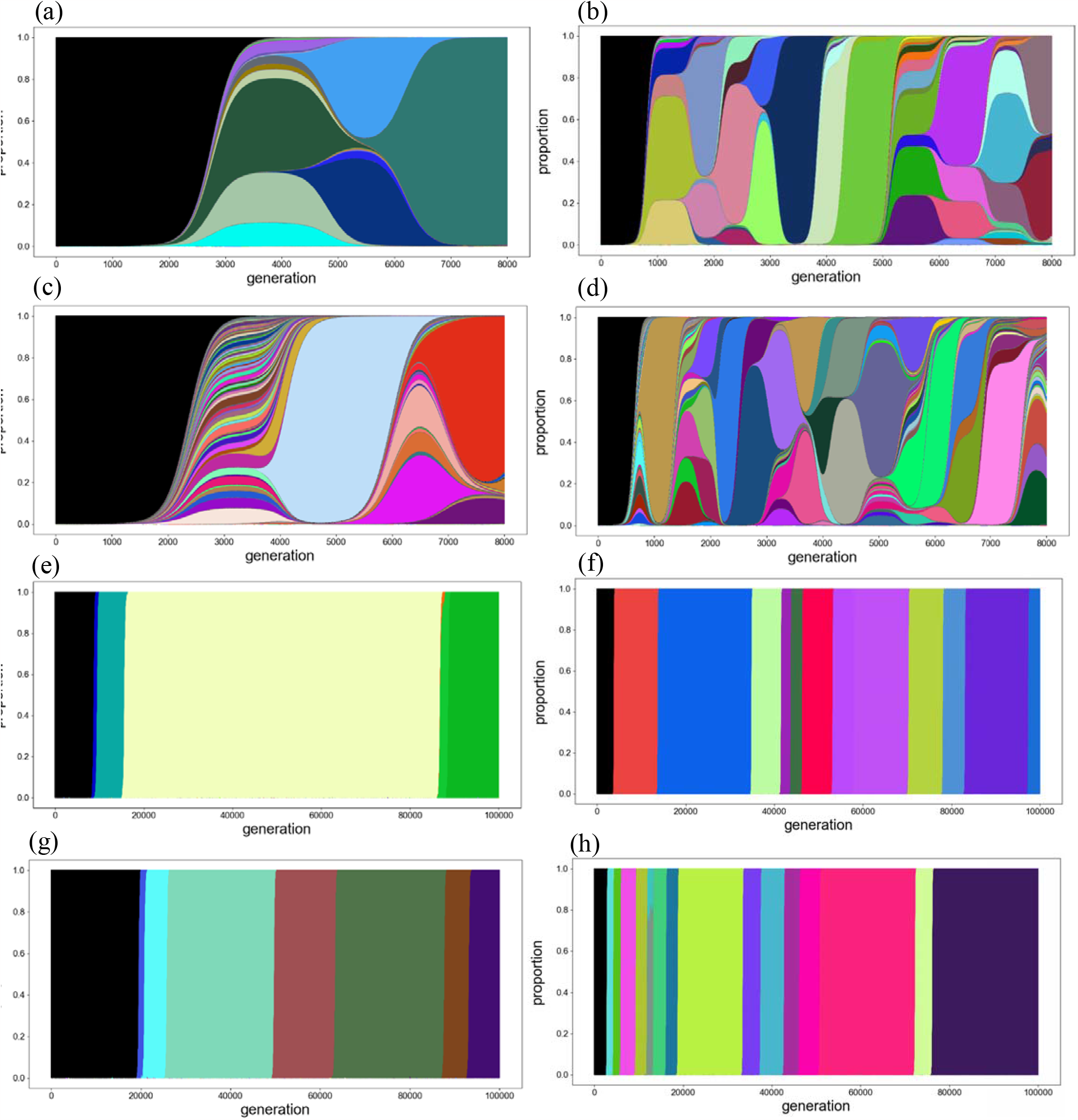
Muller plot drawn from operator modelling. The parameters in (N, m, s, generation) are (a) (10^8^, 2 × 10^-8^, 0.005, 8000), (b) (10^8^, 2 × 10^-8^, 0.02, 8000), (c) (10^8^, 2 × 10^-7^, 0.005, 8000), (d) (10^8^, 2 × 10^-7^, 0.02, 8000), (e) (10^4^, 4 × 10^-7^, 0.01, 10^5^), (f) (10^4^, 4 × 10^-7^, 0.03, 10^5^), (g) (10^4^, 8 × 10^-7^, 0.01, 10^5^), (h) (10^4^, 8 × 10^-7^, 0.03, 10^5^). From (a) to (d), clonal interference from diverse lineages concurrently occurs (concurrent regime) while from (e) to (h), one lineage sweeps the whole population (successional regime).

For the strong drift regime where N · *s*_*cr*_ ≲ 1, the simulation from operator model measured that the beneficial mutation accumulation rate *v* ≈ *α*(N, *m*, s_*cr*_) · *f* (*N*) · *m* · s_*cr*_ (for 10000 ≥N ≥50, 4× 10 ^−3^ ≥ m ≥ 10 ^−4^, 2 ×10 ^−3^ s_*cr*_ ≥10 ^−4^, which effectively suffice the N·s_*cr*_ ≲1 condition). Although there is a same chance of positive and negative s occurrence from Cauchy distribution, as Fig .2. illustrates, positive s values are more often fixed throughout the evolutionary stages, and *v* therefore tends to increase. This implies that even the small amount of selective advantage could help any mutation to be fixed in the strong drift regime. Such phenomenon becomes more apparent when N<200 as Fig. 2. (a) shows. The hypothesis for this behavior could be made from the fact that drift effect is larger for smaller N, and this means that any surviving (or, *existing, even to few*) traits from relatively larger selective benefit could be more easily amplified by chance.

**Fig. 2.**
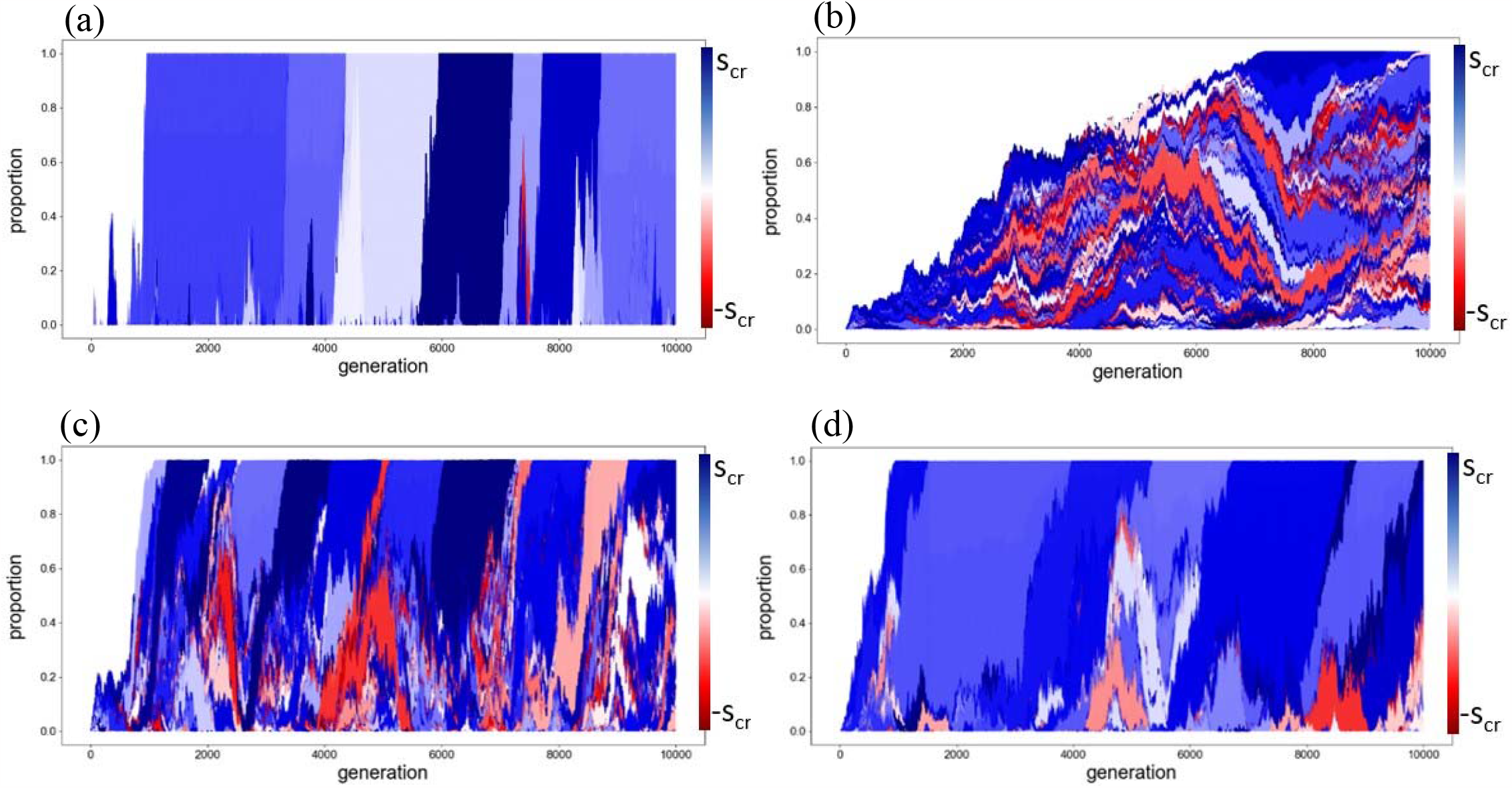
Muller plot drawn for strong drift regime. The parameters in (N, m, s_cr_, generation) representation is (a) (50, 10^-3^, 5 10^-4^, 10^4^), (b) (10^4^, 10^-3^, 5 10^-4^, 10^4^), (c) (10^3^, 4 10^-3^, 5 10^-4^, 10^4^), (d) (10^3^, 10^-3^, 2 10^-3^, 10^4^) for binary fission case. Color blue is designated for beneficial mutation and red is designated for deleterious mutation, which specifics are represented in the color bar. The drift effect becomes smaller when N gets larger (b), and positive s values are more often occurring when N gets smaller (a).

Fig.3. confirms that for any parameters within strong drift regime, the shape of the four plots for each of the N, m, s_cr_ versus *v* are all having same forms, which suggests that mathematical modelling is possible from the deterministic behaviors lying beneath strong drift regime. The linear behavior between m and *v* in Fig.3. (a) could be intuitively understood since beneficial mutations that are ultimately fixed are proportionally provided with m value. However, the linearity between s_cr_ and *v* in Fig.3. (b) may be nontrivial, since altering s_cr_ affects the shape of Cauchy distribution from which s is sampled, and more rigorous theoretical analysis would be needed to explain such linear behavior in the strong drift regime. The f(N) for Fig.3. (c) behaves like a log function with positive y intersection, and it is crudely approximated as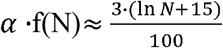. The hypothetical explanation for this logarithmic behavior could be made from two important characteristics of strong drift regime. First, as N increases, *N* · *m* increases, which provides more mutation that may increase the beneficial mutation accumulation rate. On the other hand, increased N means that probability for fixation reduces as fixation probability is nearly 1/N for small s ≈0. Then, although the mutation with relatively large s appears, such mutation may be harder to be fixed, which would hinder the beneficial mutation rate proportionally increasing with N. In sum, from the balance between increase of mutation provision and decrease of fixation, the f(N) may be logarithmically formulated. The limitation of such analysis is clear since it is grounded only on the simulation observation, and the estimated formula returns large errors for some of the data which may imply that f(N) ––––– could be far from reality and needs further modification. Nevertheless, the attempt for finding in strong drift regime demonstrated that there is a deterministic behavior behind strong stochastic (drift) effect which may be converged to –––– for some limit in the drift regime. Most importantly, such example illustrates that operator model could well provide simulation results that may lead to new findings.

**Fig. 3.**
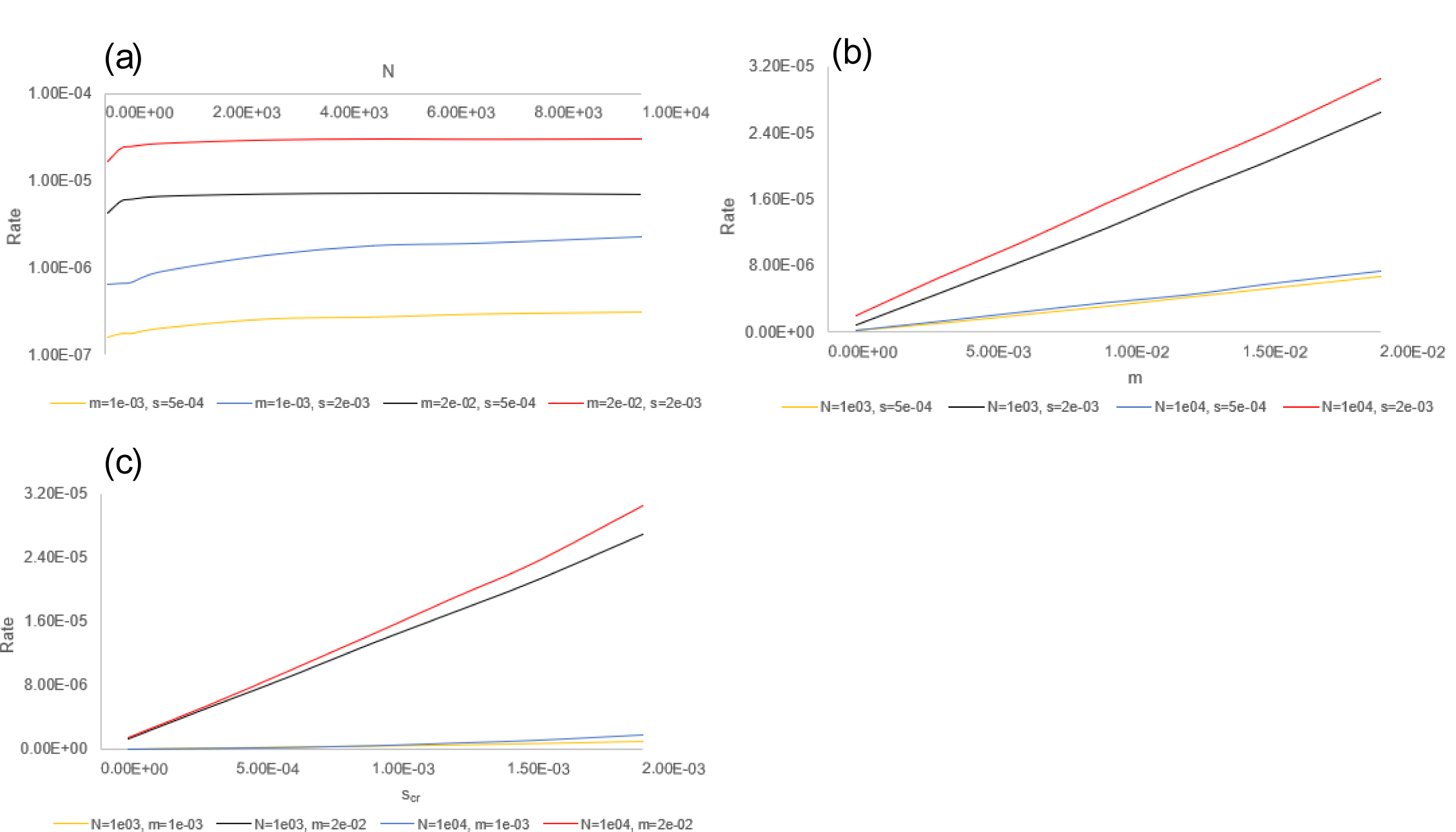
Beneficial mutation accumulation rate response with each of the N (a), m (b), s_cr_ (c), variables for strong drift regime. The parameters are set to suffice N condition. The y axis represents the rate of beneficial mutation accumulation while x axis represents each variable of interest. The y axis for (a) is represented in logarithmic scale. For each variable of N, 0.001 m 0.02, and s_cr_ 0.002, four plots were drawn to verify if the shape of plot is influenced by external variables other than the variable of interest. The four plots were chosen from the limit of minimum and maximum values of external variables of interest which is set to be 1000 and 10000 for N, 0.001 and 0.02 for m, and 0.0005 and 0.002 for s_cr_.

In nature, drift and selection exist not as distinguished processes, but rather as simultaneous stages as Moran process suggests. Therefore, distinguishing drift and selection as 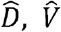 operator is, in a sense, an approximation based on the interpretation for results. Moreover, one may wonder a case if mutation operator 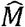 could be placed between the operation of 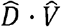 as 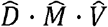 and a case where the order of drift operator and potential operator is changed as 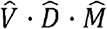. Since 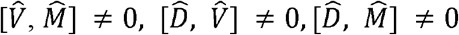 in general, such cases should be scrutinized. Here, commutator 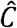 is based on the commutation relation between *Â* and 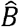 as 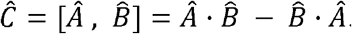.

For 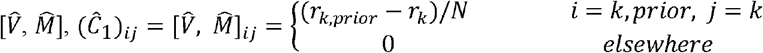 and since N ≫ 1, (*r*_*k,prior*_ − *r*_*k*_) ≪1 for most of the cases, 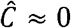. For drift operator, it is more difficult to consider the commutator since the drift operator is constructed from the prior population vector information, which would vary after selection or mutation stages. Therefore, rather than by directly computing the elements, the mean and standard deviation of the commutator for 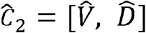 and 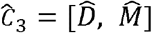 would be scrutinized. First, 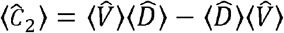 based on the assumption that all the operators are independently functioning. Now,

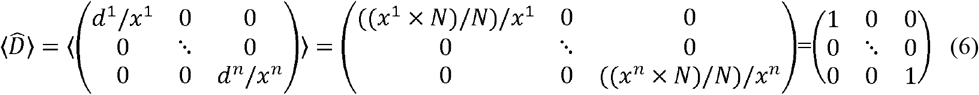

for any given population vector (Janardan and Patil, 1972). Also,

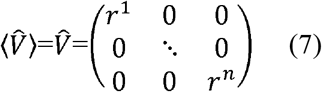

By (6) and (7), 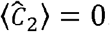. Moreover, 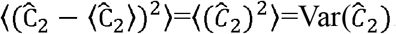. In the limit where 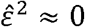, for 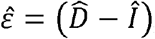, which is a realistic limit for most of the evolutionary dynamics cases (i.e. drift is not extreme), Var 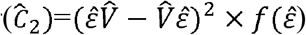where f is a probability distribution function for 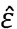. Moreover, for seldom situation where 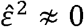 during the process of evolution, such case often has 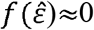. Since 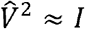 from r ≈ 1, and 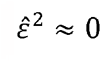 or 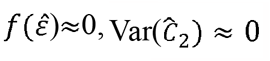 is valid for general cases where 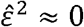 or 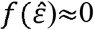 assumption is true. Thus, from the mean and variation information of 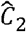, putting 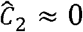for general evolutionary cases would be possible. For the 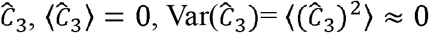 with the same reasoning for 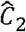 case, provided that 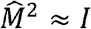, since 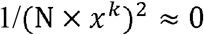 for general cases where N ×*x*^*k*^ ≫1 when new mutation occurs from kth state. Therefore, assumption that 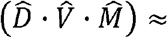 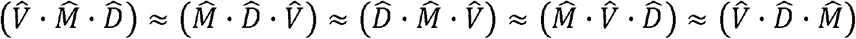 is valid for each generation. Now, the logos operator for the real world evolutionary situation, ℒ _*exact*_, would be expressed in the form of 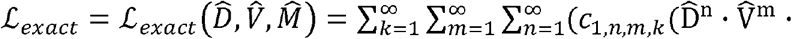 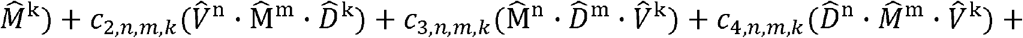 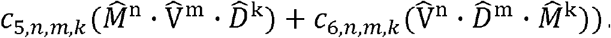.Here, *c* _*i, n,m,k*_ ≈ for *n,m,k ≥* 2 and i∈ [1,6], since terms are for the minute corrections. Moreover, to reflect that drift, mutation, and selection respectively occurred once a generation, 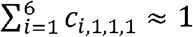. By summing all these facts, 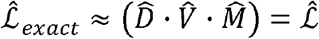 could be rendered. However, this should not be confused as 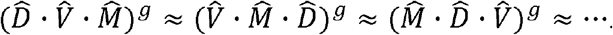 . Although for each generation, 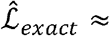 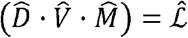, the accumulation effect of nonzero commutator values for total generation g brings about totally different stochastic results for the evolutionary dynamics when the order of 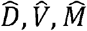 is changed. Thus, |β (g) ⟩ from 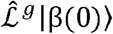 may totally differ from 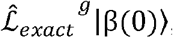, but this is far from saying that 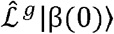 is meaningless. On the contrary, the interpretation that the operator modelling in a 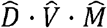 order conveys a possible picture for reality could be made. That is, among infinite possibilities of actual 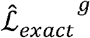process, the process where the resulting |β (t) ⟩ matches the results from 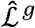 should exist, since the 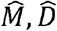 operators are in theory, able to render any population vector. Nonetheless, if the probability for the occurrence of such 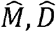 operators are too low for most of the generations, 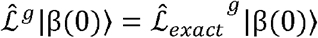may be treated only as very exceptional case. Fortunately, since 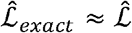 for every generation, and this implies that 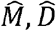 operators to match 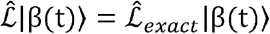 may have meaningful probability value. Therefore, the results from 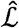 may reveal much of the information for discontinuous Markov process that could be represented as 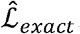. However, it should be noted that such approximation is meaningful when the commutators for the operators all approximate to 0.

Sharing the above line that 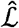 suggests one possible path from the infinitely available paths of evolutionary dynamics, the interpretation of binomial expectation for 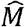 could be justified. That is, the modelling of 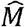 may be proceeded to cover N (total population number)-fold mutations since binomial distribution allows such minute probabilities, but since expectation of the distribution provides a sound path among diverse paths, the expectation approach could be made.

To approximate operator modelling to continuous Markov process, two methods could be introduced. The first method is defining non-integer generation values. For instance, |β (t+1/2)⟩ could be considered. The second method is allowing the overlaps of (t)⟩ with respect to generations in the form of 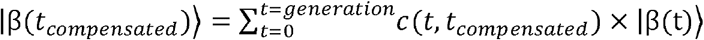 is where c (t,t compensated) is an appropriate weight factor. For advanced modelling, the first and second methods may be simultaneously used, allowing the population vector overlapping of rationally sectored generations. Specifics are introduced in the supplementary.

As the theory reveals, though the evolution is a stochastic process, such process is bounded by the non-stochastic S, E tensors. Due to their boundary-condition-like characteristics that shapes the evolutionary dynamics, these tensors are expected to render useful tools for research regarding evolutionary prediction, evolutionary control, artificial selection process, and more. Specifically, since the nomination for specific traits are possible as was done in the simulation with prime numbers (specifics in supplementary), the dynamics of nominated traits bounded by these tensors could be tracked and utilized. One suggestion for the realization of S and E tensor in experimental situation would be measuring mutation rates and fitness values in well-controlled laboratory system and conduct statistical inference for those values from large set of data. In the case where data is lacking, it may also be supplemented by the simulation with operator model, which can track the specific trait of interest, just like the experiment. Furthermore, with research regarding mutation rate and fitness landscapes, the theoretical approach for S and E tensor modelling could also be done.

Since the basic assumption is that 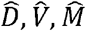 are independently operating, 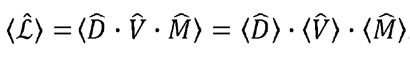 With expectation approximation of logos operator, conjecture could be made that *important relations for evolutionary dynamics* (e.g., beneficial mutation accumulation rate relation) *could be derived from such approximation* in the regime where the drift is not strong enough (i.e., N×*s* ≫ 1) thus setting 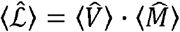 is fine. Although 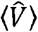 was able to be determined in the form of (6) without any further assumptions, 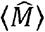 requires approximations based on the situation. With such method, the generation required for establishment and fixation were able to be found as τ_establishment_ ≈1/(*N*·*m* ·*s)* and τ _fixation_≈*ln*(*N* ·*s)/s*, which coincides with the previous results (Desai et al, 2007) which is specifically introduced in supplementary. Although the supplementary only deals with successional regime, the expectation approximation of logos may bring about important results for diverse regimes (only except for strong drift regime) as conjecture suggests.

The above process describes the evolutionary process for asexually reproducing organism. However, it could also be expected that operator modelling to cover evolutionary dynamics utilizing sexual reproduction, and such cases could be considered by distinguishing the concept of gene trait states and phenomenological trait set. Specifics are introduced in the supplementary. As the possibility of sexual reproduction operator could be discussed, if there are any other factors that involves in the evolutionary process, such factors may also be attempted to be represented as new operators. Here, the operators do not necessarily have to satisfy *linearity* as 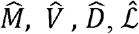 operators demonstrate. The 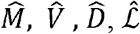 operators are *expressed* in matrix forms to act on vectors, but one must note that the essence of their operation still lies on their non-linear characteristics due to their statistical property. Moreover, as the case for the sexual reproduction operator reveals, these new operators even actually do not have to be explicitly represented as linear matrix. The sufficient condition for the new operators is, then, that these operators act on population vector and return well-defined new population vector.

## 5. Conclusion

The operator modelling is a new approach of understanding the evolutionary dynamics with expressing the evolutionary factors into intuitive form of mathematical operators, which are essentially random matrices. The simulation results showed that the new model goes along with the previous theoretical works, illustrated that general evolutionary parameters could be dealt with, and the model could lead to new findings. Moreover, the justification of the usage of operator modelling was done by commutation relation while methods to apply, reinforce, and extend the model were introduced. Such results and discussions demonstrate that operator modelling is consistent with the previous theories and is furthermore expected to provide useful insights for studying evolutionary dynamics.

## Supporting information

Simulation Data

## Availability of data and materials

All data could be found on the article, and supplementary materials.

## Code Availability

The simulation code could be found in https://github.com/Physics-physics-physics/Model.git

## Acknowledgements

We thank Professor Kyungsun Moon, Professor Su-Chan Park, and Professor Kyung Hwa Yoo for their advice on the contents and structures of the article. We also thank Changmin Park for sharing of his statistical knowledge about the hypergeometric distribution.

## Supplementary Materials

### Omission rule

The vectors, matrices, and tensors for the analysis in this article generally have infinite number of elements to reflect the infinite possibility of evolution. However, if the element is 0 for vector, or if the matrix has trivial identity part, or the index of tensor is assigned with 0 value by definition that would be introduced shortly after, the convention is set to allow those part’s omission to leave only the integral parts to be expressed. Let this convention be defined as *omission rule*. With the omission rule, the infinite element number quantities could be expressed in finite number of elements. For instance, the vector follows (A1) by omission rule convention. The 0 values could be expressed or omitted depending on its importance during analysis.

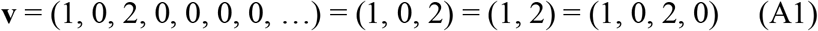

Also, matrix follow (A2) by omission rule. The trivial identity parts could be expressed or omitted, depending on its importance during analysis.

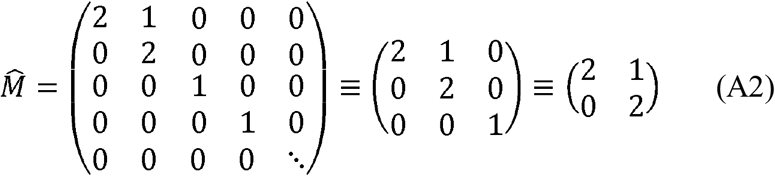

Moreover, tensor follow (A3) by omission rule.

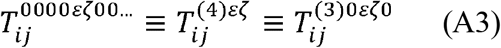

Here, the integer inside the parenthesis () for the tensor’s upper indices represents how many zeroes are present prior to the expressed indices. The representation of zeroes before the first non-zero index is important since each position of indices contains the information regarding the type of traits, which would be introduced later. With such convention in mind, index 0 for the tensor is defined to deliver the meaning that the index is not to be considered. For instance, if 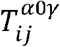, the meaningful information is determined without the consideration for β (or the second indices) and only from 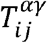. Expressing such definition in rigorous terms, the tensor 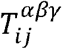 which allows the upper indices to have 0 values is equivalent to a set of 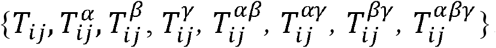. For instance, let 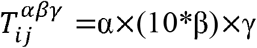 for α∈ [1,3], β∈ [1,3], γ∈ [1,3], i∈ [1,2], j∈ [1,2] and let 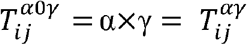 for α∈ [1,3], β∈ [1,3], i∈ [1,2], j∈ [1,2]. Then, the elements of 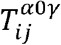 is not retrievable from the elements in 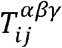. However, if the prior definition that 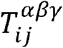is equivalent to a set that contains all the possible tensor elements using 0 indices, setting 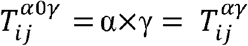is possible.

The key reason for introducing the omission rule convention is to only express the meaningful parts (which implies for the existing traits in generation t) for analysis and ignore trivial information (which implies for the possible, but non-existing traits in generation t). Hence, it should be noted that even if the quantities seem finite, or if they seem to lack some of the elements in the representations, those quantities in fact have infinite elements and unrepresented elements are already there but not expressed. Further analysis would allow more senses for this rule.

### Definition of trait

The essential assumption used in the articles was that traits could be interpreted as discontinuous quantities. In the most rigorous sense, such assumption may be invalid as height of human, for instance, is a continuous quantity. However, the assumption includes that those continuous quantities may be successfully grouped and satisfy the discontinuousness condition. For instance, the grouping of human height maybe set to be 0-100cm, 100-140cm, 140-180cm, 180-220cm, 220cm and above. Other method to apply this assumption of discontinuousness is to deal with the DNA array that correspond to phenotype of interest, since DNA sequence is essentially a discontinuous trait. For instance, population vector for DNA sequence for skin color may be expressed as P(t)={{A,T,G},{A,T,A},{A,T,C},…}. To sum up, if one may be able to correctly classify the traits with well-defined criteria, and if one may be able to track the evolution of such traits for generations, such things could be regarded as trait.

However, if the evolution of *continuous* traits that could not be discretely categorized is of major interest, and factors more than DNA sequence (e.g., environmental factors) contribute for such continuity, the continuous trait sets would have to be considered. In this case, it should not be confused that the population vector should also be a continuous quantity. The quantities regarded continuous are S, E tensors and elements in trait sets. Since the environmental capacity is set to be finite, the population vector and according 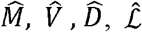 operators remain the same discontinuous form as introduced in the theory, and such discreteness is further assured by clonal interference. For the tensors, the spanning tensor would be *S* (*x,y*; *f* (*t*), *g* (*t*),*h* (*t*),…) = *m* (*x,y*), where continuous variables x, y corresponds to discontinuous integers i, j index for S tensor and f(t), g(t), … trait sets in the continuous map is from discontinuous f(t), g(t), … trait sets in the S tensor. In such continuous case, if the whole mutation rate is of interest, m_whole_ = ∬ *S* (*x,y*; *f* (*t*), *g* (*t*),*h* (*t*),…)*dxdy* could be used.

### Tensors

The current population features as well as environment plays an important role for determining the mutation rate of the population. For example, treating DNA sequence as a trait, if the trait of objective is AAAAA and if there are two current trait AAATA and CGTGC, the former would have higher mutation probability to reach the objective trait than the latter. Therefore, the current population features are integral components for determining the mutation rate for specific trait. On the other hand, as examples of temperature-conditional mutation illustrates (Tan et al, 2009), the mutation rates are often determined by the external environmental factors. The radiation, chemical mutagen, temperature, and many other environmental factors contribute to the mutation rate (Ralston, 2008). Moreover, there are lot of evidence that population density may contribute for determining the mutational rate (Krašovec et al, 2014), and mutation rate may even be a parameter for evolution itself (Zhang, 2023). Therefore, mutation rate is shaped by diverse elements, and proper reflection for interplay of these factors should be considered. For such objective, the spanning tensor S was introduced. The spanning tensor is a pre-determined (non-stochastic) map that links the population and environmental factor to mutation rate which is rendered from modelling or through the statistical analysis of experiment data in the form of S(P^k^(t_m_); environmental factor) = m^k^(t_m_) (m^k^ is a mutation rate for kth state). Although it is fine to leave S not as a tensor, the tensor representation renders advantage for sophisticated expression of mutation rates for specific traits. In this article, the external factors were set to be constant to assume that spanning tensor is a time-independent quantity during evolutionary process.

Each of the upper index of spanning tensor, which follows the omission rule, is set to correspond with each of the trait. For instance, if the first index α for *S*_*ij*_ ^*α β γ δ*…^is fixed to designate the mutation rate for trait “color” and S is constructed with such premise, S henceforth should always have the first index corresponding to the color trait. The lower indices for spanning tensor, which omission rule do not apply, designate which of the mutated traits are the interest. When the spanning tensor is constructed to have two lower indices, the first lower index corresponds to what kind of trait set the mutation is in, and second lower index corresponds to the specific element for that trait set. If one intends to classify the new mutation in different ways, the form of spanning tensor could be changed. For instance, when the interest is focused on accumulation of beneficial mutation, without any specifications of traits, the number of lower indices for spanning tensor could be set to be one and simulation was operated with such S tensor.

To thoroughly understand the spanning tensor for its usage, the above example of P^k^(T_i_) = {f_1_, g_2_} could be used. For some generation T, the mutation rate for new color f_3_ could be analyzed when f, g trait sets are given. For some biological, chemical, and physical reasons, assume that shape and original color wholly determine the mutation rate of a new color. Then, the information of the mutation rate for the f_3_ from P^k^(T_m_)= {f_1_, g_2_} could be given from *S* _13_^12^. Furthermore, there is a possibility for the mutation of independent trait set, h (wing existence) which is totally new from f, g, may occur for generation T. In this case, generation higher than T would also have to account for the h trait set for the states P^k^(t T+1) (k ∈ [1,*n*]) for analysis. Furthermore, the mutation rate for such h trait would be determined by *S*_31_^12^.

It would be meaningful to note that spanning tensor could be used for diverse objectives. For instance, since the f_3_ mutation could happen elsewhere from the kth state of generation T, the probability as a whole (i.e., for all states) for the f_3_ to occur in generation T is given as (A4).

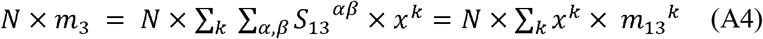

### Mutation for pre-existing traits

Although the case where the mutation for pre-existing traits was not dealt with in the article, it could be treated with two methods. One is setting upper right off-diagonal terms for the mutation operator, and the other is recollecting the |P(T) ⟩ information after mutation operation and combining the population vector elements with the same traits to retain the mutation matrix form to only have lower left off-diagonal terms. Both methods share same meaning. Let {{AT}, {AG}} be the original population for generation t, where |P(T) ⟩=(x^1^, x^2^)^T^=(0.6, 0.4)^T^, and N=100. Now, assume that binomial expectation for {AG} to {AT} mutation be given with 0.1. Then, with the probability of 0.1, the 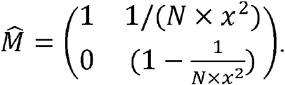 Same situation with the second method, the 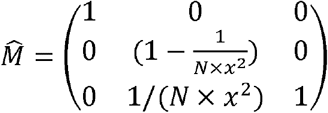 with probability 0.1, but after |P(T) ⟩ went through mutation to have (0.6, 0.39, 0.01)^T^ vector, operation could be set to add 0.01 element to 0.6 element to give the resulting (0.61, 0.39)^T^ vector.

### Set representation

First, the set representation defines a map *φ* which acts on each state P^k^(t) (k ∈ [1,*n(t)*]) and returns the vector of form 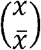 where 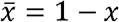, and *x* is the frequency of that state in generation t. For example, 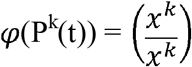, and since x^k^ value signifies the frequency for state k, 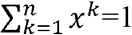. Note that, 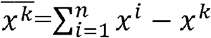is a trivial result. For the priorly introduced example of P^k^(T) = {f_1_, g_2_}, when Hardy-Weinberg equilibrium is met (Edwards, 2008) after generation T, the *φ*(P^k^(t))(k ∈ [1, *n*]) would have constant elements for the vectors for t≥T, thus n(t)=n(T) for t ≥T. However, when the equilibrium is broken, the evolution comes into play and elements of each vector may change as the generation goes on. Moreover, even the new trait elements f_3_, g_3_, h_1_ could be added to the trait set f, g, h if any new mutations occur and change n(t). Now, from the vector formed by map *φ*, the mutation could be expressed as an operator that is essentially a random matrix [random matrix ref] that depends on the current population vector. Assume that the total population number is N(t) for each generation t. For generation T, the color f_3_ would be mutated from the state P^k^(T_m_) with the rate of *N* × *x*^*k*^× *m*_13_^*k*^, and mutation operator 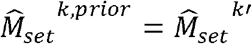 would have a form of

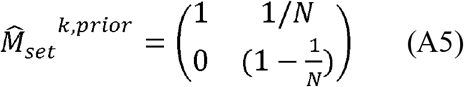

that acts on the 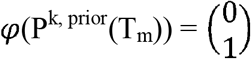. That is, when mutation occurs for T_m_, the currently non-existing trait of f_3_ is *priorly* prepared from the T_m_ generation set. Here, after P^k, prior^(T_m_) is *realized*, it should be designated with specific k’th state, but since P^k, prior^(T_m_) is not an existing state yet, it would be left as *k, prior* state to suggest for its yet non-determined characteristics. Moreover, if 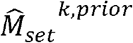 is not an identity matrix, so does for 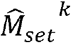 which has a form of

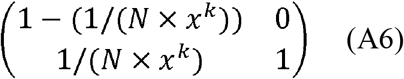

that acts on 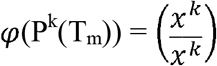 Therefore, the resulting P(T_v_) would contain the k’th state P^k’^(T_v_)={f_3_, g_2_} as its element where 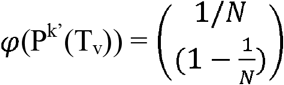 and kth state P^k^(T_v_)={f_1_, g_2_} where 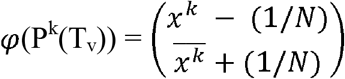. If the mutation does not occur by the rate of (1− *N* × *x*^*k*^× *m*_13_^*k*^ for the kth state, the mutation matrices acting on the vectors would be the identity matrix of size 2 × 2 for both the *φ* (P^k, prior^(T_m_)) and *φ* (P^k^(T_m_)). Note that among all T stages, only Tm stage should consider the *prior* elements.

Regarding the above example of P^k^(T_m_) = {f_1_, g_2_}, the random mutation matrices for 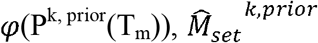 could be given as (A5) with possible P^k, prior^(T_m_) and its mutation probability. When 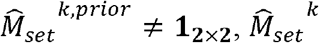 is automatically (A6). In sum, the mutation operators that contribute to new trait k’ from k state are always acting pairwise to each of the *φ* (P^k, prior^(T_m_)) and *φ* (P^k^(T_m_)).

After the mutation step, the population faces selective pressure. To express this process in the set representation, the potential operator has the form of

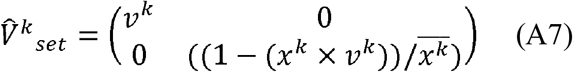

that acts on 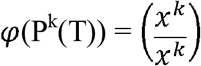, and *v*^*k*^ = (*x*^*k*^ × *r*^*k*^)/*weight*, where *E*^*α β γ δ*…^ = *r*^*k*^ and weight 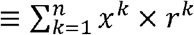.

With the example of P^k^(T_v_) = {f_1_, g_2_},

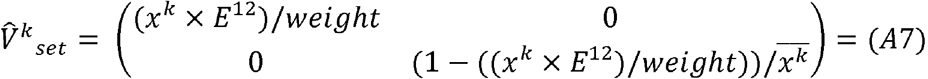. The *φ* (P^k^(T)= {f_1_, g_2_}) after the selective pressure would be given as 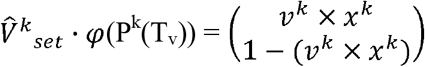 and still preserve the form 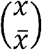.Such analysis could be done for ∀k [1, n(T)].

To represent the result of drift in set representation, with (*u*^*k*^/*N*) ≡ *d*^*k*^,

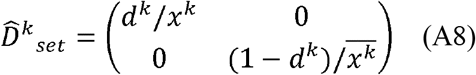

(A8) acts on 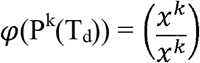 to return 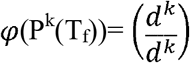. Here, one could note that the information of *x*^*k*^ is already included in the process of getting *d*^*k*^ values. Moreover,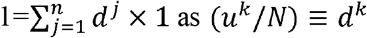.

The logos operator for the set representation 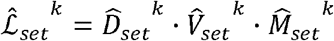 would have the form of 2 × 2 matrix with possibly off-diagonal term from mutation operator. Then,

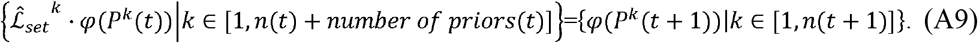

The n(t) here signifies the number of states in P(t) set, and number of priors(t) represents the total possible number of mutation traits from all of the current states. Here, similar to the omission rule convention, if 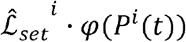 returns 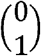 which means that the element *P*^*i*^ (*t* + 1)is actually non-existing, such element could be omitted in the set { *φ P*^*k*^ (*t* + 1)) | *k* ∈ [1,*n* (*t* +1)]}. Analyzing such set representation result throughout the generation, the derivation of P(t+1) information from that of P(t) is possible.

The set representation clarifies the details of evolutionary process and reduces the operator size, but it conveys relatively less intuitive picture. Moreover, the tensor product approximation is unavailable in this approach. On the other hand, the whole representation conveys more direct picture of what is going on in the evolution process for generation T than the set representation, with tensor product approximation also being possible. Nevertheless, the size of 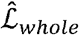 matrix may be too large if 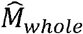has too many elements to consider for the large number of priors(t) and eventually become unwieldly to utilize for the computation. In short, both representations convey the same meaning, and proper choice of representation for the objective of analysis would be recommended.

### Tensor direct product approximation

There is one point where the whole representation has advantage over the set representation. That is, when the distinct traits are almost independently contributing for the determination of mutation rate. The map *ζ* that acts upon the trait set element should be introduced for further explanation. Specifically, *ζ* satisfies the relation for above trait f, g as *ζ* (f_i_ (T)) = X_fi_(T), *ζ* (g_i_ (T)) = X_gi_(T) where x is the frequency of that trait and 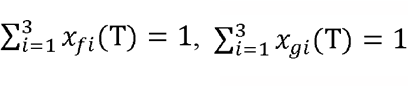. Moreover, trait set vectors |f (T) ⟩,|g (T) ⟩ is defined in a way that |f (T) ⟩_*i*_ ≡ *ζ* (f_i_(T)), |g(T)⟩_*i*_ ≡ *ζ* g_i_ (T)) . With the map *ζ*, the detail process of mutation for P^k^(T_m_) ={f_1_(T_m_), g_2_(T_m_)} could be analyzed. Since the traits are almost independently contributing to the mutation rate, *ζ* (f_1_) × *ζ (g*_*2*_)= *ψ* (P^k^(T_m_)) + O(P^k^ (Tm)) *ψ* ≅ (P^k^(T_m_)). This would be applied to ∀ k ∈[1, n], which would lead to the interpretation that

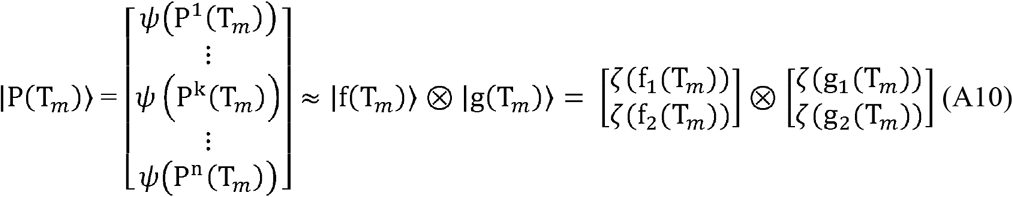

for the whole representation. In this case, since mutation rate is almost independently determined by the distinctive traits (i.e. f, g), the mutation rate of generation T could be given from spanning tensor as 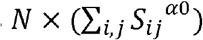 for trait f_α_’s mutation, 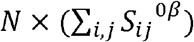 for trait g_β_’s mutation. With these rates, mutation operators for |f (T) ⟩,|g (T) ⟩ could be constructed in a way which could directly act upon each of the 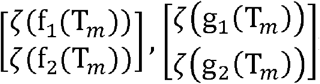 rather than on the

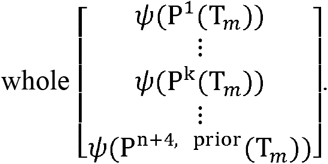

For details, the prior element for the trait |f (T*m*) ⟩,|g (T*m*) ⟩ would be expressed as

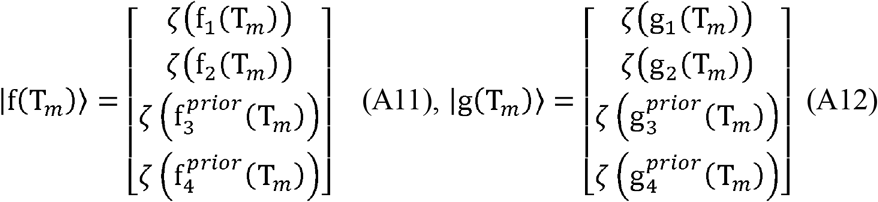

where

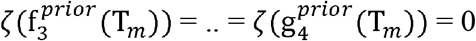. Combining the results, the mutation operator for |P (T*m*) ⟩ could be constructed as Table A1.

**Table A1.**
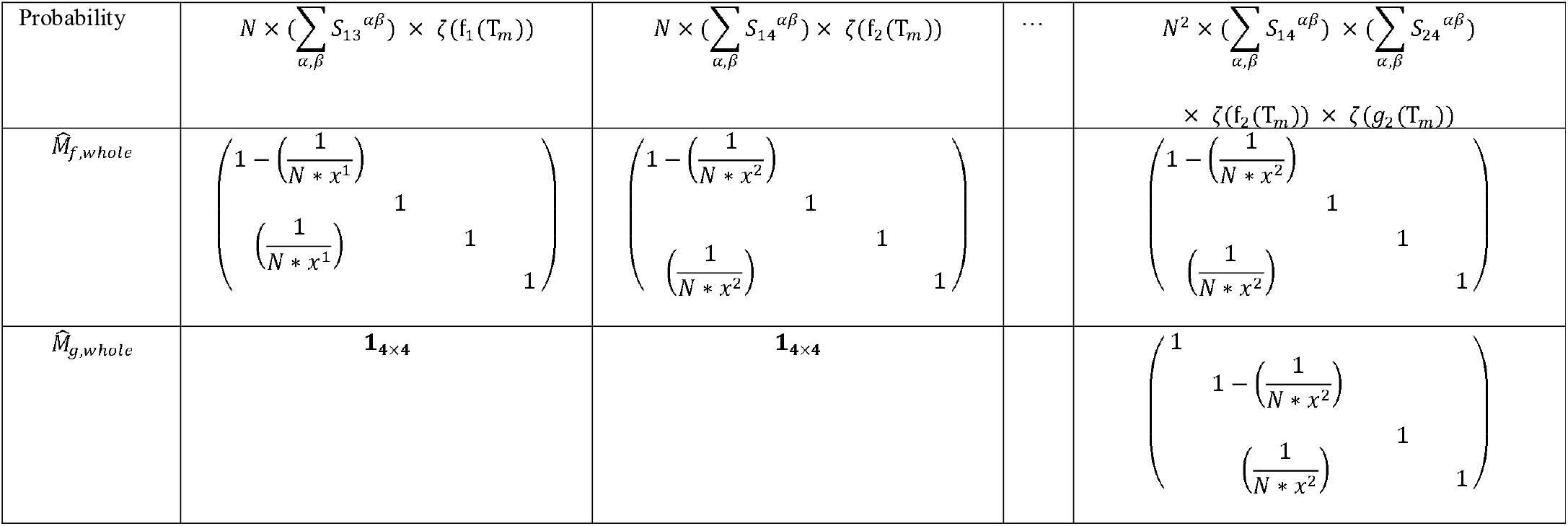
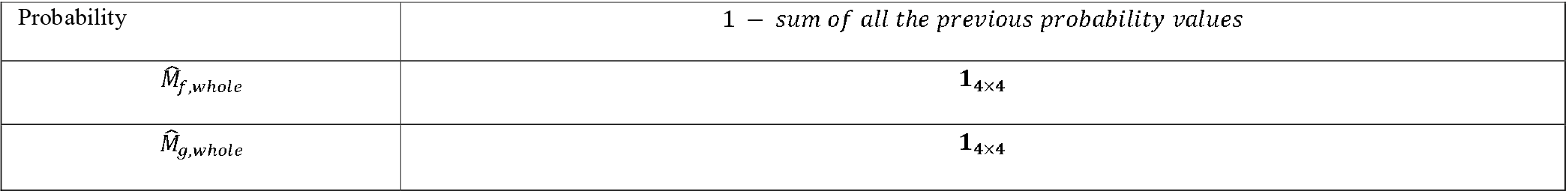
Random matrix representations for trait set vectors.

Now, the whole mutation operator in this case would be approximated as 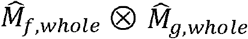 with corresponding mutation rate. However, 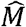 does not have to be fully computed since the tensor computation allow the following:

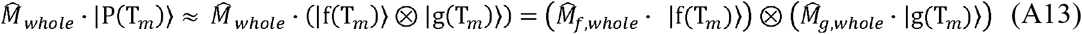

Here, |f (T*m*) ⟩,|g (T*m*) ⟩ satisfies (A11), (A12).

For such tensor approximation case, if the new trait set h is introduced by allowing the new mutated trait element *h*_*1*_, extra consideration of 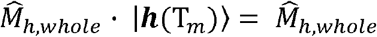 · 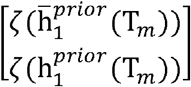should be made. Here, 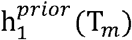corresponds to the prior trait of generation T and 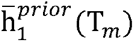corresponds to the prior trait of generation T which is the complementary element of 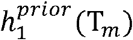. It is natural to put 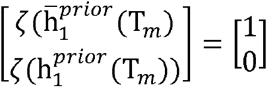 when the meaning of 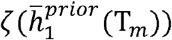 is the complement of the frequency for the population that has trait 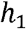 for the generation T. To such vector, 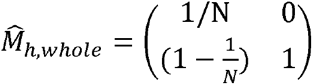 is acted upon with specific mutation rate that depends on the original proportion of traits. Lastly, in the limit where the traits are almost independently contributing to determining the fitness, *ζ* (f_1_) × *ζ (g*_*2*_)= *ψ* (P^k^(T_v_)) + O(P^k^ (T_v_)) *ψ* ≅ (P^k^(T_v_)), the potential operator for f, g trait would be 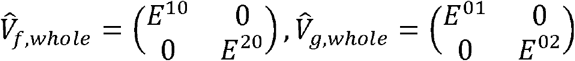 and these operators would act on 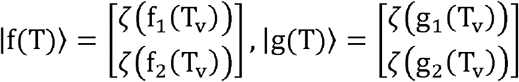 individually. Therefore,

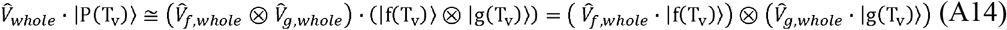

Since the elements of resulting vector for 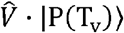 may not sum up to 1 in such tensor approximation case, the weight factor should divide the vector for final |P (T) ⟩ Lastly, it must be noted that condition of *ζ* (f_1_) (T_v_))) × *ζ (g*_*2*_) (T_v_)) ≅ *ψ* (P^k^(T_v_)) for selection stage and condition of *ζ* (f_1_) (T_m_)) × *ζ (g*_*2*_) (T_m_)) ≅*ψ* ≅ (P^k^(T_m_)) for mutation stage are not the same. For instance, the mutation rate may largely depend on the interdependent interactions between currently existing traits, thus tensor product approximation for the mutation stage may not be valid. Nevertheless, the interaction between the traits for determining the selection strength may be almost independent and tensor product approximation could be well utilized in such case. Therefore, the tensor approximation method could be used for potential stage, while its application to mutation stage would not be possible.

For the case of mutation and selection, the tensor product approximation was possible. However, for the drift, since the operator deals with the total P(t) and distinction of trait sets in this case is meaningless, tensor product approximation method, which primarily focuses on the separation of |P (t) ⟩ for distinct traits, would not be introduced.

### Expectation approximation of logos

The expectation approximation is used to reduce the degree of freedom for the multivariate functions. To apply such approximation in the operator model, let the consideration be done for the situation of simulation where 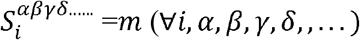, and *E*^*α β γ δ*……^ =*s* × (number of nonzero indices *α β γ δ…*.). Here, the subtlety between the traits is not considered, and each new mutations are regarded as a single element of the new trait set. To see how the expectation approximation for logos works, let initial |P (t) ⟩ ≡ (*x*^1^.*x*^2^,0,0,…)^T^, . For such population vector, the expectation of mutation operator could be represented as the following.

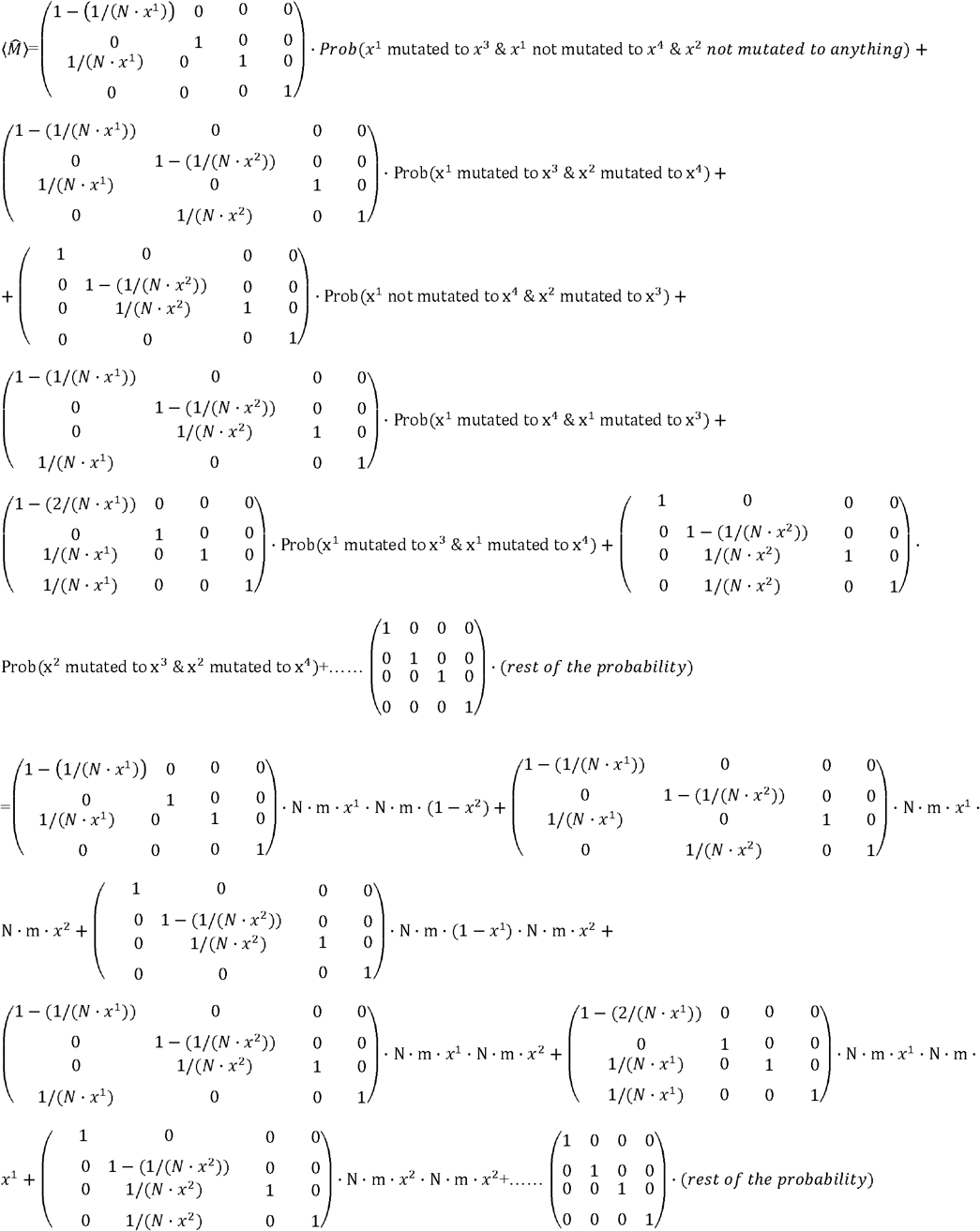

is the mean of mutation operator. Theoretically, such expansion could go to infinity since x^5^, x^6^, …. could be infinitely considered. However, for now cases, let N ·*m <1* thus N·*m*·*x* < 1 and (N·*m*·*x)*^*2*^ ≪1, so the power of 2 and above could be safely ignored. In such situation,

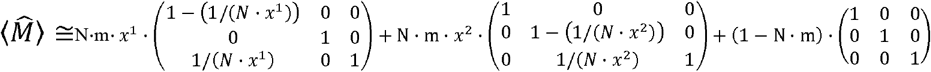

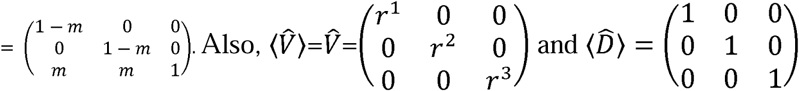

Therefore, in the limit where (N ·*m* ·*x*)^2^ ≪ 1, with the assumption that 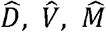 are independently acting on the population which is robustly discussed above, 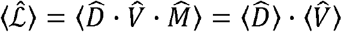 .

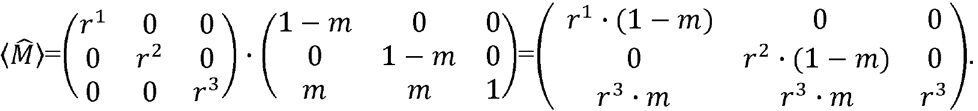

As could be seen here, the expectation of logos operator produces matrix that do not depend on the proportion of the current population vector elements, but only on N, m, and s (r=1+s). With appropriate approximations, the expectation of logos operator is expected to render useful results in many ways, since these matrices are only composed with parameters for evolutionary dynamics. To briefly get a sense for the application of expectation approximation of logos operator, the derivation process of generation required for establishment and fixation of new beneficial mutation would be introduced. To do this, assume a situation where only beneficial mutations are occurring, |P (0)⟩ ≡ (1, 0, 0, 0, …)^T^, the dynamics is occurring at successional regime where N ·*m* ≪1, and fitness is determined only from the number of accumulated beneficial mutations. Setting *r*^1^=1 and *r*^2^=r=1+s, the mean logos operator would have the form of 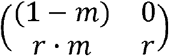Moreover, since the elimination of mutations from the drift should be reflected on 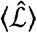 though 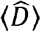is not suit for such objective, the components represented on 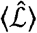 should be designed to represent only those who survived the drift, which in other words are established and should show deterministic actions. With such fact, let there be assumption that there is no extra mutation prior to first mutation state reaching the proportion of 1/ (*N* ·*S*), which is the proportion where the state would be established.

Now, since the new mutation occurs with the probability of N · *m*, the average period for the occurrence of new mutation would be 1/ (*N* · *m*). By assumption, since there is no additional mutation until the point of the new mutation’s established, for 1/ (N · *m*) generations, 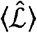 would have the form of 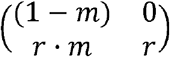. Then, from the assumption that *m*^*k*^ ≈ 0for k≥0,

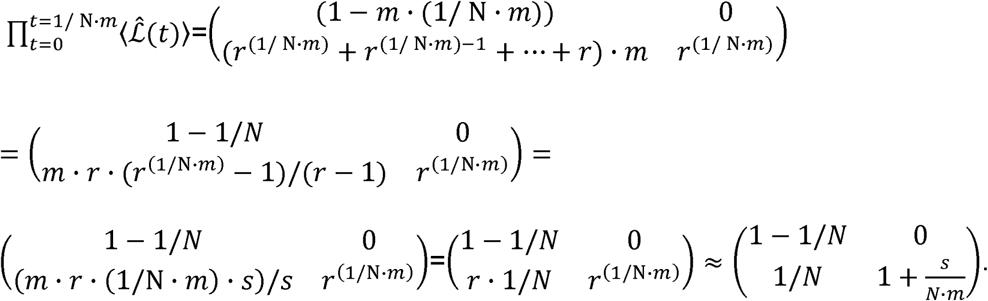

By acting |P (0) ⟩ upon 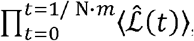, the result is 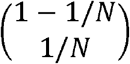, which is expected. Now, to find the generation for establishment, let such generation be τ. Then, with the given assumptions,

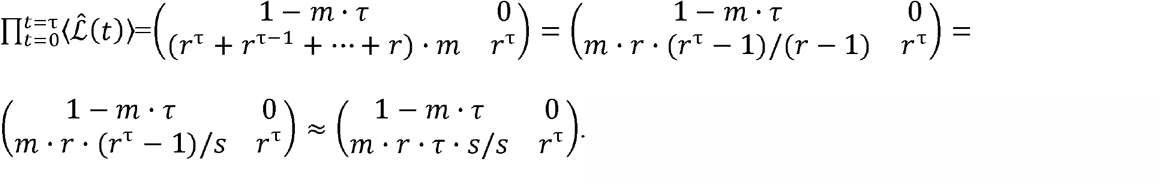

By acting such operator to |P (0) ⟩, the expected result is 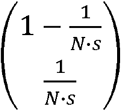 . Therefore,*r* · *τ* ·*S*/(*S*·*m*) ≈ 1/ *(S*·*N)*., and this leads to ≈ 1/ *(N*· *m*·*S)*, which goes along with the previous result. It should be noted that the weight was put to 1 here, since the 1− *m* · *τ* ≈1and *m* · *r* · *τ* ≈0. Using the result of *τ*≈ 1/ *(N*· *m*·*S* the beneficial mutation accumulation rate v=s/*τ N*· *m*·*s*^*2*^ for such successional regime.

Now, for the generation taken for fixation, the form of logos expectation is expanded, since new second mutation could now be provided, hence having the form of 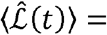 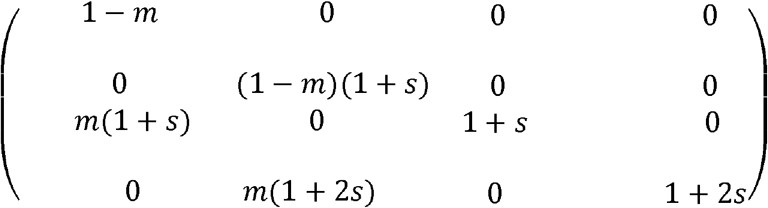 for t ≳1 *N*· *m*·*S* in the setting that x^3^ is from x^1^ and x^4^ is from x^2^. However, since the analysis is focused on the fixation of first mutation and other mutations are out of interest, let the natural assumption that 1-m ≈ 1be applied, and 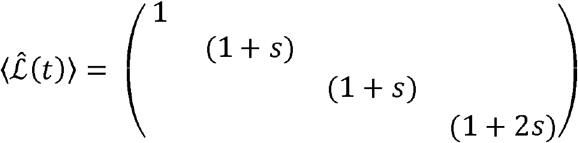 . This matrix would act on the operator 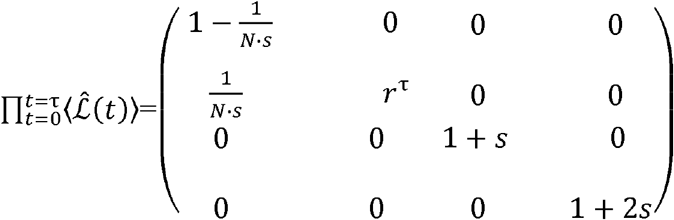 . Since the interested part in the expectation of logos here is the upper left 2× matrix part, only those part would be scrutinized. Let the time for fixation be τ’. Then, 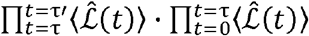 gives the result of

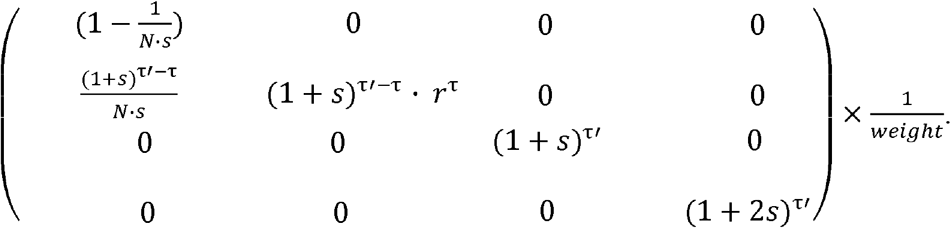

If such operator acts on |P (0)⟩, it renders the result of 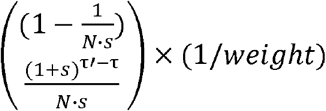. Here, since 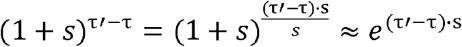 . Important part here then, is the comparison between the two elements 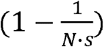 and 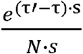 since weight 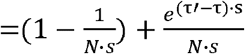 and if (second element/weight) = 1/2, it could be roughly interpreted that the second element caught up with the original first element, and thus the second element is fixed. This leads to 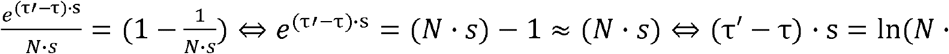 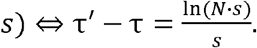. Therefore, the time for fixation is approximately in the scale of 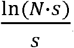. Although there were many assumptions for rendering the results, the conjecture is that far general analysis for evolutionary dynamics could be made with expectation of logos operator, if any sound assumptions and interpretations for the situation is given.

### Continuous Markov

The operator method may try to reflect characteristics of continuous Markov model by two adjustments. The first adjustment is defining non-integer, rational number generation between discrete integer generation. For instance, between t and t+1, the t+1/2 could be defined, and if portion of population begins the reproducing process (e.g., for bacteria, mitosis initiation) closer to t+1/2 rather than t, those population may be expressed in |P (t + ½) ⟩ rather than |P (t) ⟩ . From such scale of generation separation by half, the amount of computation process would be increasing by at most 2^2^. This is because the population vector size would approximately be doubled, and generation to consider, in effect, is also doubled. For example, assume that integer generation population was initially defined to be P(*t* _*integer number*_) = {….., {(f_1,_ g_2_)(t)….}. Now, if t and t+1/2 are separated, P(*t* _*rational number*_) = {….., {f_1,_ g_2_} (t),{f_1,_ g_2_} (t +1/2),….}= P(*t* _*rational number*_+1/2) and total amount of generation iteration to consider increased from t to 2t.

For the operators, the 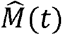 operator may contain non-diagonal element 1/N only if the generation for population vector matches the generation for the mutation operator. To rigorously express this in mathematics, 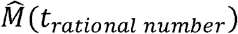 and 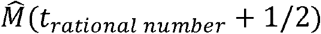 operators only contain non-diagonal terms when they are respectively operating to |P (t_*rational number*_) ⟩ and |P (t_*rational number*_+1/2) ⟩ The 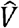 operator could also be distinguished for t and t+1/2. Specifically, 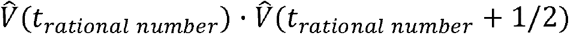 should be equal to 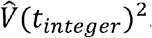. Since 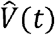 is derived from environment tensors which have values independent of generation, and since 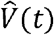 is a diagonal matrix, the diagonal components of 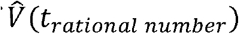 are square root values of diagonal values of 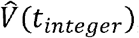, by the relation 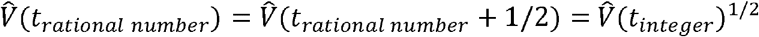.

The 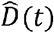 is more difficult to deal with, because of its essentially stochastic property. There are three possible considerations for constructing 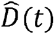. The first one is to let 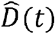 be distinguished for t and t+1/2 as 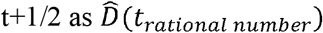 and 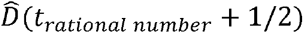. This is to say that drift operator only acts on the vector components of their generation and leave the rest of the components untouched. The second one is to let 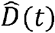 be operating for the whole population vector of rational number generations only at the integer generation. The last one is to let 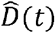 be operating on every sectioned rational generation for the whole population vector, with modifying (3) into

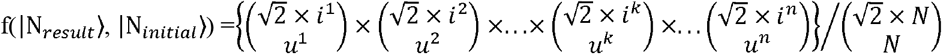

where 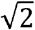 is from the assumption that during half generation, the population may theoretically increase 2^1/2^ fold when there is no pressure from environmental capacity.

Since the drift operator is an approximation of drift phenomenon, all three methods are in a sense providing meaningful information in the macroscopic viewpoint with their own advantages and shortcomings. The first model distinguishes the moments when certain drift operator should operate, and such model is especially useful when cell cycle has lot of influence for the drift phenomena. That is, if there are any case where asexual organisms die off or survive more easily during some specific point of their cell cycle, the effect of drift could be weakened or strengthened based on in which of the cell cycle the population is going through, and such consideration could be done by the first model. In fact, by comparing with the experimental results, the coefficient 2 for *i*′s in (20) is expected to be adjusted in this model. This is because the population part that is set to be non-drift-experiencing for certain moment, in reality, is interacting with the drift-experiencing part of the population, and such phenomena should be considered. This method, however, has major drawback. That is, if the population portion for t and t+1/2 differs too much so that one part of portion is too small, the multivariate hypergeometric distribution constructed with such information may not properly reflect the reality. Setting aside the quantitative analysis for such phenomenon, the qualitative explanation could be made by the example of P (t) ={….,{f_1_, g_2_}(t) {f_1_, g_2_}(t + 1/2),…..} = *P* (*t* + 1/2) . In the example, one should note that P(t+1/2) and P(t), although they share same trait, would be interpreted as different states. Moreover, let *ψ* ({f_1_, g_2_}(t)) = 0.30 and *ψ* ({f_1_, g_2_}(t + 1/2)) = 0.04, and assume that 0.05 is the portion for establishment of the population. In reality, {f_1_, g_2_}would not be influenced much from the drift, since the total portion value of 0.34 represents that such state went far beyond the establishment step, and the portion would be less likely to decrease drastically. However, the {f_1_, g_2_}(t + 1/2) could easily be influenced by drift, thus the total population of 0.34 in this case is vulnerable to large portion of decrease. To sum up, to use the first method, the proportions for P^k^(t) or P^k^(t+1/2) for certain state should not be too small. The second method, which lets the drift operator to act only on the integer periods of the population, may not properly reflect the differences of drift phenomena for P(t) and P(t+1/2) in the expense of decreasing the amount of computation and avoiding the problems of first method. The third method tries to reflect the drift happening for states in both P(t) and P(t+1/2), which is most like realistic case. However, the seemingly intuitive assumption of letting the coefficient of *i* of multivariate hypergeometric distribution to be 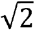may be naïve and needs further correction. Specifically, let one generation be sectored to 1000 fine, rational number generations, and binary fission is taking place. In such case, the coefficient *i* for multivariate hypergeometric distribution would be 2^1/1000^ ≈1 and cannot properly depict the situation for drift. Therefore, correction of coefficient based on the experiment and theory is expected to render higher reliability for the third method. If such correction could be properly provided, the third method is expected to bring best precision among three methods suggested. Lastly, it should be noted once again if generation interval is sectored n times, the operator model may attain the characteristics of continuous Markov process accordingly, but the amount of computation would at most increase by n^2^.

Now, the second adjustment could be done by allowing the population vectors of contiguous generations to be overlapped by pre-determined weight factors. In detail, 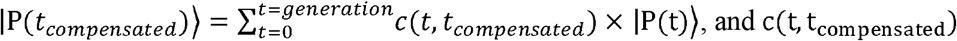, and C (t,t _compensated_)which is a weight representing the extent of overlapping of population defined for contiguous generations, could be determined from the experimental results and knowledge about cell cycle of the organism. For instance, |P (*T*_*compensated*_) ⟩=0.05 × |P (T−2) ⟩ + 0.15 × |P (T− 1) + 0.6 × |P (T) ⟩ × + 0.15× |P (T+1) ⟩ +0.05× |P (T+2) form of modelling may be possible.

For advanced modelling, the first and second adjustments may be simultaneously used, allowing the overlapping of population vectors that are defined from rationally sectored generations. However, it should be noted that though the first and second adjustments to meet the characteristics of continuous Markov process may work out, they are still an approximation for the real, continuous Markov process cases.

The operator modelling may differ from reality if each asexual organism’s reproduction period deviates too much from one another, and thus may also require increased adjustments to compensate for such variances. However, there are some possible ways in laboratory to control the deviation of cell cycle, with *cell synchronization* methods (Chen and Deng, 2018; Banfalvi, 2011). With such methods, the deviation of the reproduction rate may be controlled and provide conditions where operator method could be well utilized.

### Specific information for the simulation

It should be noted that although the set representation was used in the actual codes, the following accounts are made with whole representation, which is simpler to explain with. One of the most important parts of simulation was to store the lineage information of mutations. To do this, prime numbers were used. That is, when two new mutations were provided from preexisting trait corresponding to p1*p2*p3, the new mutations were respectively allocated with p1*p2*p3*p4 and p1*p2*p3*p5. Since multiplication results of different prime numbers are never the same, all the mutations were able to be distinguished, and the lineage information could also be stored in such way. Now, to consider for the mutation rate, the constant distribution between [0,1] was used. In detail, if mutation rate was set to be m, and any random number x between [0,1] chosen from constant distribution sufficed x<m, the prior term for population vector was filled with 1/N when N× m<1. When N × m>1, the population vector was provided with 1/N to the [N × m] prior terms, and when the randomly chosen value from constant distribution was less than N × m-[N × m], the last 1/N was added to the population vector. Many of the case, the initial population was set to be P(0)={{b0}} and corresponding vector be |P (0) ⟩ ≡ (*x*_1_-1, 0, 0, …)^T^. However, the codes were also made to deal with the situation where initial population have more than one trait. Now, if the element of |P (0) ⟩ becomes less than 1/N by drift or selection process, the algorithm was set to give 0 for such element. Repeating such process for given total generations, the information for evolutionary dynamics process was given. For each combination of parameters of (N (total population number), s (fitness increase), m (mutation rate), g (generation), i (initial mutation varieties)),300 iterations (or 100 iterations for some of the strong drift regime simulations) were made and proportion of simulation results and theoretical results for fixation rate and mutation accumulation rate were measured. With these 300 (or 100) values for each of the proportion of fixation rate and mutation accumulation rate, the null hypothesis was that the histogram from values would follow Gaussian distribution with mean value 1, and KS test was done to find out if such hypothesis is correct. Among diverse candidates, number of bins for the histograms from 300 (or 100) values were chosen to maximize the p-value for KS test. Moreover, when alpha parameters had to be estimated for measuring fixation rate for concurrent regime, mean value of alpha values from first 50 iterations were found. To verify that such mean value is the correct estimation, the rest of the 250 iterations used such mean value to see if the histogram follows Gaussian distribution with mean value around 1. The simulation was also made to deal with dynamical population case. To consider such case, especially when the population is increasing, the Gompertz model of bacterial growth was introduced in the form of *N* (*t*) = *N*_0_ · *exp* (*ln* (*N* ∞ */N*_0_). *exp* (− *exp* (*μe*·(λ ∞*t)/* (*ln* (*N* ∞ */N*_0_)+1))) where *N*_0_ is initial population number, *N* ∞ is environmental capacity, is the maximal value of the first derivative for growth curve, λ is the lag time (Zwietering et al, 1990). For the simulation of dynamical population case, especially in the log phase of the growth curve, the drift was assumed to be low since offspring would be less likely to diminish when the population did not meet environmental capacity. To model such situation, the coefficient of multivariate hypergeometric function was adjusted with sigmoid function having N(t)/*N*_0_ as its domain value.

### Sexual reproduction operator

To consider for the sexual reproduction operator, two trait sets, which is *gene trait set* and corresponding *phenomenological trait set* should also be defined. For instance, example gene trait with the above P^k^(t), the phenomenological {*f*_1_, *g*_2_}state could be represented into diploid gene traits state {{(*f*_1_), (*f*_1_)},{ (*g*_2_), (*g*_2_)}}. Furthermore, if gene traits for f and g are linked, {{(*f*_1,_ *g*_2_), (*f*_1,_ *g*_2_)}} classification could be done. Now, the mutation operator acts on gene trait while the potential operator acts on phenomenological states. It should be noted that, if *f*_1_ is a dominant trait whereas *f*_2_ is a recessive trait, both gene states {{(*f*_1_), (*f*_2_)},{ (*g*_2_), (*g*_2_)}}, {{(*f*_1_), (*f*_2_)},{ (*g*_2_), (*g*_2_)}} corresponds to the same phenomenological state of {*f*_1,_ *g*_2_}. However, although the phenomenological traits seem alike, they should be distinguished as different states with same trait set elements, since the underlying gene states are different. It should be noted that as well as the mutation operator, the drift operator should also act on the gene traits. For the mutation and potential operators, the spanning and environmental tensor could still be applied in the same way as the asexual reproduction case. Lastly, the sexual reproduction operator acts on the gene trait which acts on population vector and produces new vector. Thus, sexual reproduction operator may, in theory, be represented as linear matrix form as it is operating as a linear map between two vectors. However, since the representation process for the sexual reproduction operator is cumbersome while the detailed form of the matrix is not necessarily important for finding the result, the specific form of sexual reproduction operator would not be introduced for now. Rather, the resulting vector after the operation of sexual operator would be introduced since this is a major part of interest. Let the gene trait for generation t be composed of P(t) = {{{(*f*_1_), (*f*_2_)},{ (*g*_2_), (*g*_2_)}}, {{(*f*_1_), (*f*_1_)},{ (*g*_2_), (*g*_2_)}}}and corresponding |P (t) ⟩ be |P (*t*_*i*_) ⟩.= (0.8,0.2,0,…)^*T*^ Then, the gene pool for *p*_*f*1_ = (0.8/2) + (0.2) = 0.6, *p*_*f*2_ =0.8/2 =0.4 *p*_*g*2_ =1. Therefore, for the offspring, P(t+1)_initial_= {{{(*f*_1_), (*f*_1_)},{ (*g*_2_) (*g*_2_)}}, {{(*f*_1_), (*f*_2_)},{ (*g*_2_) (*g*_2_)}}, {{(*f*_2_), (*f*_2_)},{ (*g*_2_) (*g*_2_)}}} and corresponding |P(t+1)_initial_ = ((0.6^2^),2 × (0.6 × 0.4)^2^)^*T*^ which components adds up to 1. As could be found here, the population vector for the analysis of sexual reproduction deals with larger data than the previous asexual case, and amount of computation is larger. Modelling with such sexual reproduction operator is expected to return useful information which could enlighten new perspectives towards studying Muller’s rachet (Muller, 1964; Andersson and Hughes, 1996), red queen hypothesis (Strotz et al, 2018) and more.

## References

Aalto, E. (1989). The Moran model and validity of the diffusion approximation in population genetics. Journal of theoretical biology, 140(3), 317–326.

Andersson, D. I., & Hughes, D. (1996). Muller’s ratchet decreases fitness of a DNA-based microbe. Proceedings of the National Academy of Sciences, 93(2), 906–907.

Baake, E., & Gabriel, W. (2000). Biological evolution through mutation, selection, and drift: An introductory review. Annual Reviews of Computational Physics, 7, 203–264.

Banfalvi, G. (2011). Overview of cell synchronization. Cell cycle synchronization: methods and protocols, 1–23.

Bräutigam, C., & Smerlak, M. (2022). Diffusion approximations in population genetics and the rate of Muller’s ratchet. Journal of Theoretical Biology, 550, 111236.

Chen, G., & Deng, X. (2018). Cell synchronization by double thymidine block. Bio-protocol, 8(17), e2994–e2994.

Chesson, J. (1976). A non-central multivariate hypergeometric distribution arising from biased sampling with application to selective predation. Journal of Applied Probability, 13(4), 795–797.

De Oliveira, L. R. (2014). Master Equation: Biological Applications and Thermodynamic Description.

Desai, M. M., Fisher, D. S., & Murray, A. W. (2007). The speed of evolution and maintenance of variation in asexual populations. Current biology, 17(5), 385–394.

De Visser, J. A. G., & Krug, J. (2014). Empirical fitness landscapes and the predictability of evolution. Nature Reviews Genetics, 15(7), 480–490.

Edwards, A. W. F. (2008). GH Hardy (1908) and hardy–Weinberg equilibrium. Genetics,179(3), 1143–1150.

Ewens, W. J. (2004). Mathematical population genetics: theoretical introduction (Vol. 27, pp. xx+-417). New York: Springer.

Fog, A. (2005). Simulation models for biological and cultural evolution. Socially Inspired Computing, 21.

Fog, A. (2008). Calculation methods for Wallenius’ noncentral hypergeometric distribution. Communications in Statistics—Simulation and Computation®, 37(2), 258–273.

García-Dorado, A., Avila, V., Sanchez-Molano, E., Manrique, A., & Lopez-Fanjul, C. (2007). The build up of mutation–selection–drift balance in laboratory Drosophila populations. Evolution, 61(3), 653–665.

Gavrilets, S. (2010). High-dimensional fitness landscapes and speciation. Evolution: the extended synthesis, 45–79.

Gerrish, P. J., & Lenski, R. E. (1998). The fate of competing beneficial mutations in an asexual population. Genetica, 102, 127–144.

Janardan, K. G., & Patil, G. P. (1972). A unified approach for a class of multivariate hypergeometric models. Sankhyā: The Indian Journal of Statistics, Series A, 363–376.

Krašovec, R., Belavkin, R. V., Aston, J. A., Channon, A., Aston, E., Rash, B. M., … & Knight, C. G. (2014). Mutation rate plasticity in rifampicin resistance depends on Escherichia coli cell–cell interactions. Nature communications, 5(1), 3742.

Levy, S. F., Blundell, J. R., Venkataram, S., Petrov, D. A., Fisher, D. S., & Sherlock, G. (2015). Quantitative evolutionary dynamics using high-resolution lineage tracking. Nature, 519(7542), 181–186.

Miralles, R., Gerrish, P. J., Moya, A., & Elena, S. F. (1999). Clonal interference and the evolution of RNA viruses. Science, 285(5434), 1745–1747.

Moran, P. A. P. (1958, January). Random processes in genetics. In Mathematical proceedings of the cambridge philosophical society (Vol. 54, No. 1, pp. 60–71). Cambridge University Press.

Muirhead, C. A., & Wakeley, J. (2009). Modeling multiallelic selection using a Moran model. Genetics, 182(4), 1141–1157.

Muller, H. J. (1964). The relation of recombination to mutational advance. Mutation Research/Fundamental and Molecular Mechanisms of Mutagenesis, 1(1), 2–9.

Nguyen Ba, A. N., Cvijović, I., Rojas Echenique, J. I., Lawrence, K. R., Rego-Costa, A., Liu, X., … & Desai, M. M. (2019). High-resolution lineage tracking reveals travelling wave of adaptation in laboratory yeast. Nature, 575(7783), 494–499.

Park, S. C., & Krug, J. (2007). Clonal interference in large populations. Proceedings of the National Academy of Sciences, 104(46), 18135–18140.

Ralston, A. (2008). Environmental mutagens, cell signalling and DNA repair. Nature Education, 1(1), 114.

Strotz, L. C., Simoes, M., Girard, M. G., Breitkreuz, L., Kimmig, J., & Lieberman, B. S. (2018). Getting somewhere with the Red Queen: chasing a biologically modern definition of the hypothesis. Biology letters, 14(5), 20170734.

Tan, G., Chen, M., Foote, C., & Tan, C. (2009). Temperature-sensitive mutations made easy: generating conditional mutations by using temperature-sensitive inteins that function within different temperature ranges. Genetics, 183(1), 13–22.

Tran, T. D., Hofrichter, J., & Jost, J. (2013). An introduction to the mathematical structure of the Wright–Fisher model of population genetics. Theory in Biosciences, 132(2), 73–82.

Wakeley, J. (2005). The limits of theoretical population genetics. Genetics, 169(1), 1–7.

Walsh, B., & Lynch, M. (2018). Evolution and selection of quantitative traits. Oxford University Press.

Wright, S. (1949). The genetical structure of populations. Annals of eugenics, 15(1), 323–354.

Zhang, G. (2023). The mutation rate as an evolving trait. Nature Reviews Genetics, 24(1), 3–3.

Zwietering, M. H., Jongenburger, I., Rombouts, F. M., & Van’t Riet, K. J. A. E. M. (1990). Modeling of the bacterial growth curve. Applied and environmental microbiology, 56(6), 1875–1881.

